# Differential DNA methylation in the *Vinv* promoter region controls Cold Induced Sweetening in potato

**DOI:** 10.1101/2020.04.26.062562

**Authors:** L. Shumbe, M. Visse, E. Soares, I. Smit, B. Dupuis, H. Vanderschuren

## Abstract

Control of potato sprouting is important to ensure constant supply of high-quality potato to the industry. Efficient control of sprouting can be achieved by chemical treatment or cold temperature. Recent bans on anti-sprouting molecules are prompting the use of cold storage in the potato value chain. Unfortunately, storage of potato at low temperatures is associated with cold induced sweetening (CIS) due to the induction of the vacuolar invertase gene under low temperatures. Because CIS is associated with the production of the potentially carcinogenic acrylamide in processed potatoes, concise knowledge on the regulatory mechanisms controlling the CIS-phenotype in potatoes is expected to help pave the way for the production of CIS-resistant potato varieties. Here, we dissect the promoters of the Vacuolar invertase (*Vinv*) genes from CIS-susceptible and CIS-resistant varieties to investigate their implication in CIS-phenotype determination. Using bisulfite sequencing and CRISPR-dCas9-DRM2-mediated *de novo* DNA methylation, we show that the CIS-resistant phenotype of Verdi, is in part due to hypermethylation of its *Vinv* promoter, more specifically in the 1.0-1.7kb region. Those findings open new perspectives to engineer CIS-resistant potatoes by genome and epigenome modifications.

## Introduction

A major problem faced by the potato processing industry nowadays is finding a balance between sprout control during storage and quality of the processed products. The recent ban on the use of Chlorpropham (CIPC) for sprout control in the European union (EU2019/989 of 17 June 2019) results from its potentially toxic and carcinogenic effects^1,2^ and the detection of CIPC residues and its primary metabolite 3-chloroaniline in raw and processed potatoes^3,4^. In this context there is an urgent need to develop efficient and safe alternatives to CIPC to control sprouting in the potato value chain.

Alternatives such as ethylene^5^, 1, 4-Dimethylnapthalene (1,4-DMN)^6^, 3-Decen-2-one^7^ and Maleic hydrazide^8^ have been used over the years in the EU, USA and Canada. However, compounds such as 1,4-DMN are also subjected to Maximum Residue Level (MRL) in the EU, implying a degree of unsafety. Safer alternatives are provided by natural essential oils such as Spearmint oil (BIOX-M), Caraway oil and Clove oil^9–11^. However, most of these natural oils require high concentrations to achieve effective sprout inhibition, thus rendering them less cost effective.

The use of anti-sprouting chemicals can be circumvented by storing potatoes at physiologically inactive temperature. This cost-effective approach also minimizes water loss and reduces disease incidence^12^. However, low temperature storage at 4 °C enhances the breakdown of starch to reducing sugars (glucose and fructose) leading to the so-called Cold induced Sweetening (CIS)^13^. Upon processing at high temperatures, these reducing sugars undergo a complex reaction with amino acids, specifically asparagine, to form brown to black colored products in the Maillard reaction^14^. Acrylamide, a toxic and potentially carcinogenic compound has been identified as one of the colored products generated during the Maillard reaction and a strong correlation has been established between chips/fries color and acrylamide content^15,16^.

Starch is metabolized to reducing sugars by a series of enzymes catalyzing the reactions of starch breakdown and hexogenesis. Several studies have demonstrated the implication of these processes in CIS^16–20^. However, the amount of reducing sugars generated in the potato tubers is the function of a synergistic action of the enzymes catalyzing the afore-mentioned processes and the interrelated pathways of starch synthesis, glycolysis, and respiration^13,21^. The bottleneck for CIS is the conversion of sucrose to glucose and fructose, catalyzed mainly by invertases and sucrose synthase^13,21^. A strong correlation between vacuolar invertase (VINV) activity and reducing sugars content makes *Vinv* the primary determinant of CIS in potato tubers^16,22,23^. In addition to its regulation by cold temperatures, the VINV enzymatic activity is also controlled *in planta* by inhibitors (INH)^22,24^. Brummell and co-workers identified INH1 and INH2 as the two most abundant forms of invertase inhibitors in the potato genotypes 1021/1 and 937/3 and localized INH1 in the apoplast and INH2 in the vacuole, with *INH2α* allelic form coding for the full length INH2 protein^25^.

The importance of CIS in potato varieties with good processing properties and the need to reduce the use of toxic sprouting inhibitors such as CIPC is prompting the potato value chain to opt for varieties such as Verdi, Lady Claire, Multa and White Pearl with reported low accumulation of reducing sugars over long term storage at 4 °C^26–28^. The availability of genotypes with CIS-resistant phenotype offers opportunities to investigate the genetic basis for low CIS in potato under cold conditions. In this study, commercial potato varieties contrasting for CIS were used to investigate the genetic and epigenetic contributions to CIS.

## Results

### Reducing sugars determine acrylamide content of cold stored processed potato

Acrylamide formation through the Maillard reaction requires two principal precursors: reducing sugars and amino acids, specifically asparagine^29^. Accumulation of reducing sugars in potato tubers is a function of breakdown rates of starch to sucrose and sucrose to glucose and fructose. Assessment of total reducing sugars (glucose + fructose) in tubers after 1, 3 and 5 months of storage at 4° C and 9° C allowed classifying the potato varieties (var.) according to their reducing sugar levels: high (var. Nico), average (var. Bintje and var. Lady Rosetta (Laro)) and low (var. Verdi) (Fig. 1A). Based on these levels, the potato varieties could be classified into CIS-resistant variety (Verdi) and CIS-susceptible varieties (Bintje, Nico and Laro). Our results also confirmed that tubers stored at 4° C for all CIS-susceptible varieties accumulate significantly higher amounts of reducing sugars relative to their counterparts stored at 9° C^27^. Noticeably, reducing sugars accumulated with time of storage for Bintje, Nico and Laro, but decreased with storage time in Verdi. These trends inversely correlated with the sucrose contents in Bintje, Nico and Laro which showed a general trend of reduction in the sucrose content with time of storage whereas sucrose levels in Verdi tubers increased during storage (Fig. 1B).

**Fig. 1:**
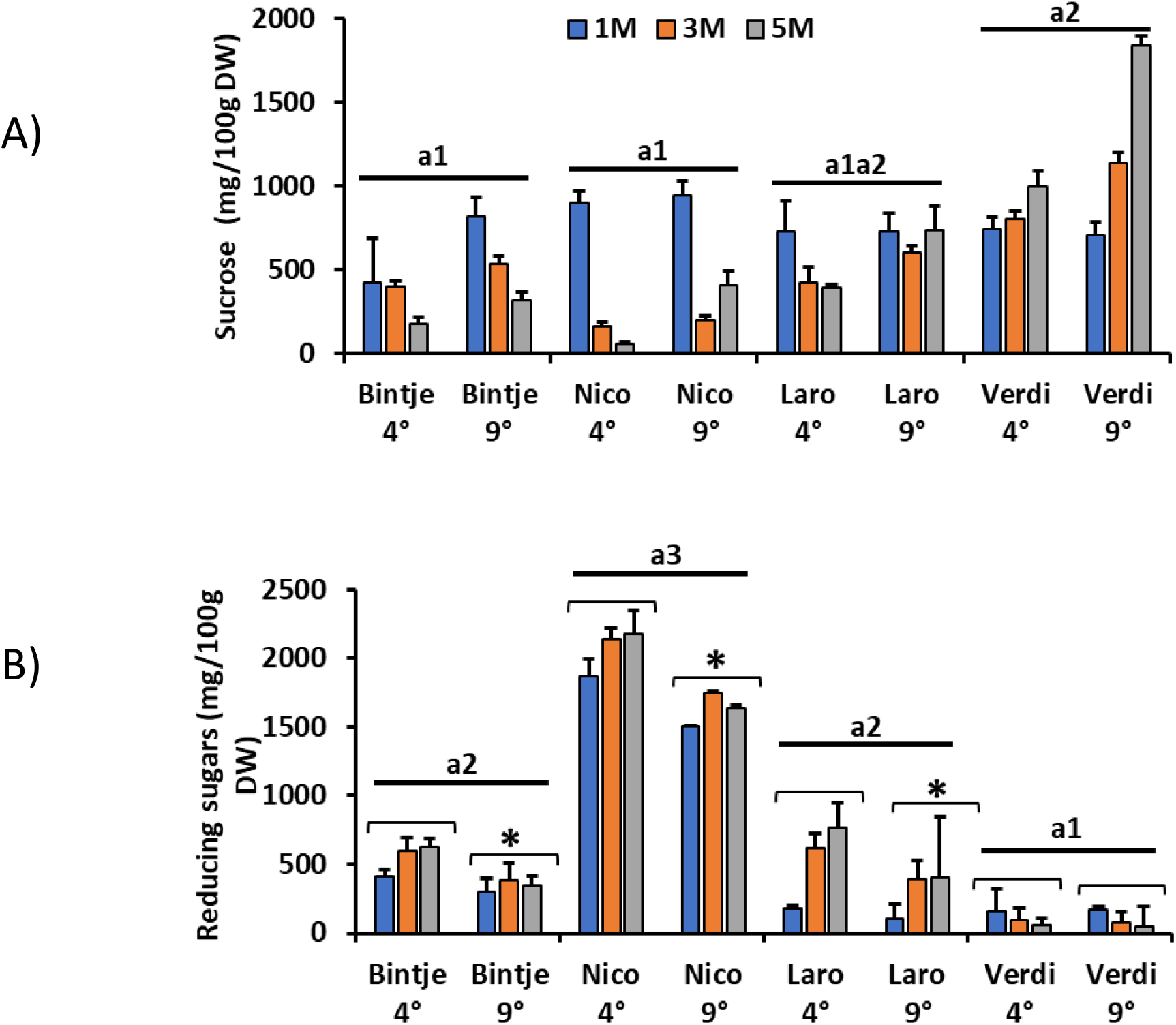
CIS-phenotype of potato varieties after 5-months cold storage. **A)** Sucrose content and **B)** content of reducing sugars (glucose + fructose) assayed from freeze-dried tubers of Bintje, Nico, Laro and Verdi after 1, 3 and 5 months of storage at 4° C and 9° C. n =4-5 independent biological replicates. a1,a2,a3 represent statistically significant differences between varieties at P<0.05 computed by Tukey test and * represents significant differences between storage temperatures for each variety at p<0.05 computed using the student’s t-test.

We also assessed the effect of low temperature storage on the content of free asparagine in the 3 month-stored potato varieties (4° C and 9° C). Our results revealed that storage temperature had no significant effect on the level of free asparagine for all the varieties (Fig. 2A). However, the level of free asparagine in Bintje was significantly lower than that in Nico, Laro and Verdi, indicative of a variety-dependent regulation of free asparagine content. The temperature-dependent regulation of reducing sugars suggests that in potato varieties accumulating similar amounts of asparagine, reducing sugars are the limiting factor for acrylamide formation upon cold storage and processing. We processed two commercial chips varieties Laro and Verdi displaying contrasting levels of reducing sugars (Fig. 1B) and similar levels of asparagine (Fig. 2A) after 5-month storage at 4 °C and controlled CO_2_. The chips produced from Laro tubers were remarkably dark brown to black in color, compared with the clear chips obtained from Verdi tubers (Fig. 2B(ii)). The color of the potato chips consistently correlated with acrylamide content (acrylamide content of 10413 μg/kg and 319 μg/kg in Laro and Verdi chips respectively, Fig. 2B(iii)), and glucose content (Fig. 2B(i)). These results corroborates previous findings that potato chips color is proportionate to acrylamide content^15^, making color of chips a reliable qualitative measure of the level of acrylamide.

**Fig. 2:**
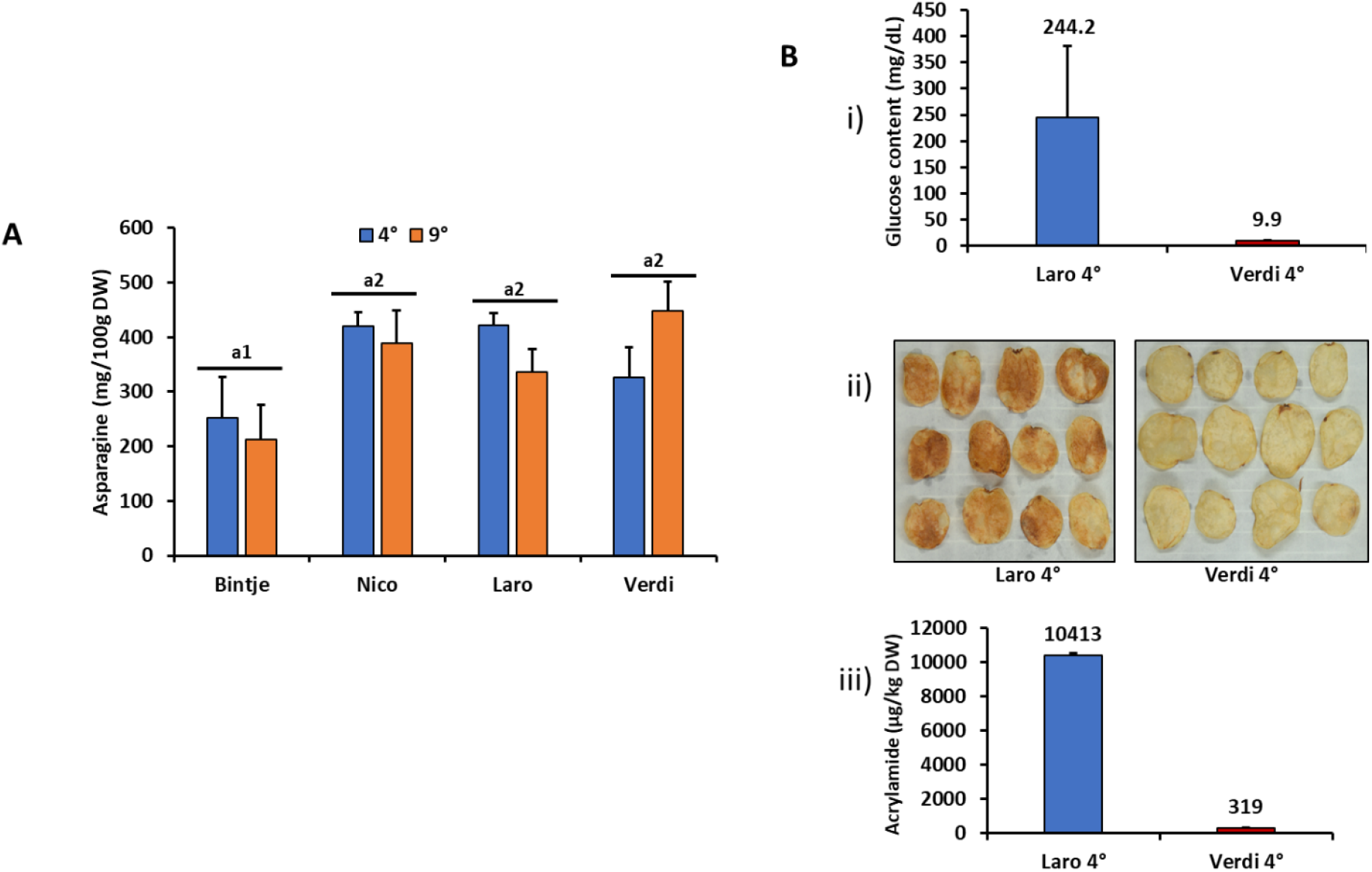
Effect of cold storage on asparagine and acrylamide content. **A)** Asparagine content from Bintje, Nico, Laro and Verdi potato tubers after 3 months of storage at 4° C and 9° C, n=5. a1, a2, a3 represent statistically significant differences between varieties at P<0.05, computed using the Tukey test. **B)** Glucose content (**i**) of Laro and Verdi tubers after 5-months storage at 4° C and controlled CO_2_ measured using a glucometer diabetic strip (Accu Check), n=3. Glucose values of 9.9 mg/dl represent the minimum detection limit of the glucometer (**ii**) corresponding quality of processed potato chips assessed by chips color and (**iii**) acrylamide content of technical replicates obtained from two independent extractions from the chips (**ii**).

### VINV and invertase inhibitors control CIS in potato

Several studies have positioned VINV as the bottle-neck for CIS, based on the correlation between its gene expression (and/or protein activity) and the levels of reducing sugars^16,27^. *Vinv* gene expression was examined in the potato tubers after 1, 3 and 5 months of storage at 4° C and 9° C by RT-qPCR. Transcript levels of *Vinv* in tubers stored at 4° C were generally higher than in their corresponding varieties stored at 9 °C (Fig. 3A). This observation is in line with the levels of reducing sugars at the respective temperatures (Fig 1B). In like manner, the general transcript levels of *Vinv* in the CIS-susceptible varieties Nico and Laro were significantly higher than the levels in the CIS-resistant variety Verdi. Noticeably, Bintje and Verdi had similar *Vinv* transcript levels. An inverse correlation was observed between the transcript levels of *Vinv* in Verdi and its level of reducing sugars: *Vinv* transcript levels increased slightly with time of storage whereas reducing sugar levels decreased with time of storage (Fig. 1B and 3A). Although globally there exists a correlation between the levels of *Vinv* in Nico, Laro and Verdi and their respective reducing sugar levels, this correlation is not perfect as evidenced by the above-mentioned discrepancies in Bintje and Verdi. Recent studies have shown that the regulation of VINV activity is also controlled at the post-translation level by acid invertase inhibitors in potato and other plant models like sugarcane^22,25,30^. Therefore, we decided to assay the activity of VINV in the selected potato varieties. Because *Vinv* transcript levels after 1 and 3 months of storage in the selected potato varieties showed good correlation with the levels of reducing sugars, we decided to assay the acid invertase activity from tuber samples stored for 3 months at 4° C. The acid invertase activity from Bintje tubers was significantly higher than that from Verdi tubers (Fig. 3B), contrary to the similar *Vinv* transcript levels observed for these varieties (Fig. 3A). Highest activity was recorded for Nico, with Bintje and Laro showing significantly higher activities relative to Verdi (Fig. 3B). The activity measurements highly correlated with the respective reducing sugars levels for all tested varieties.

**Fig. 3:**
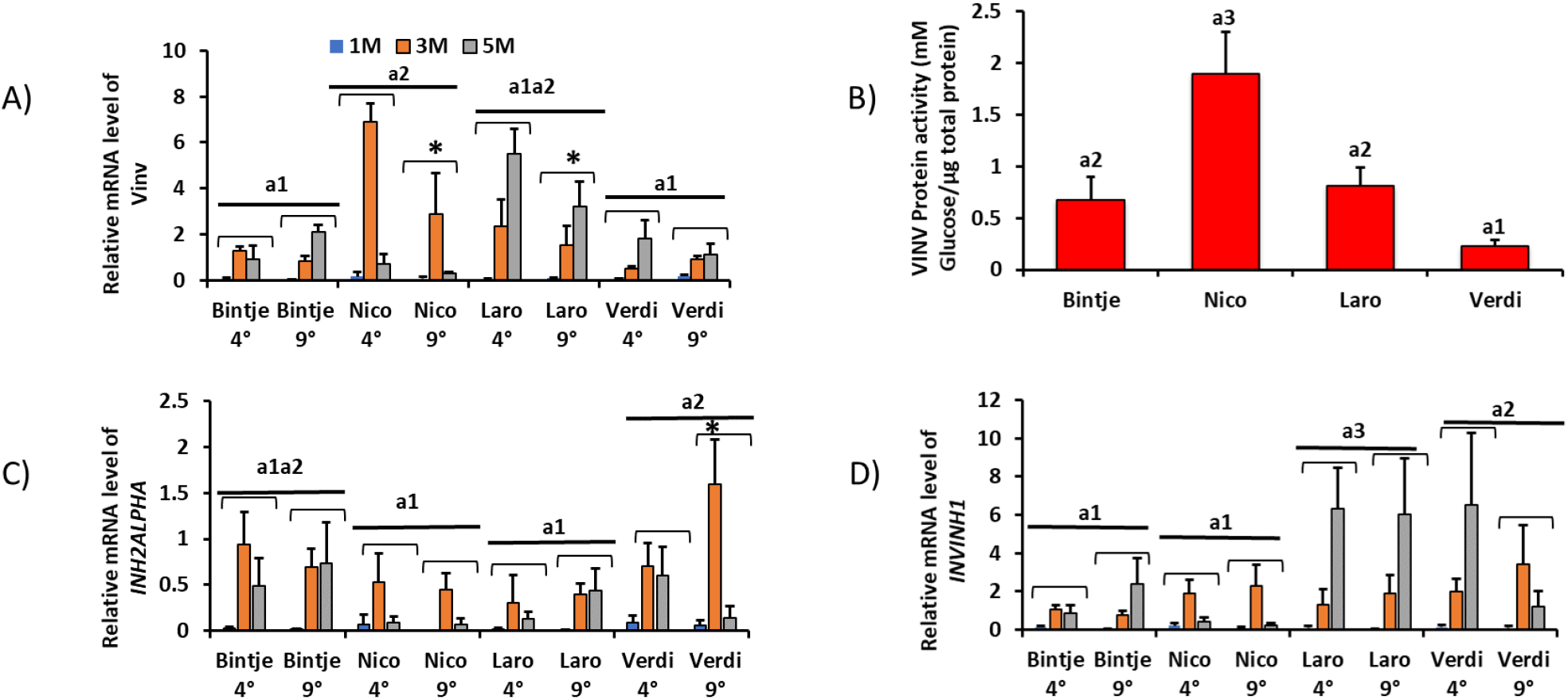
Role of vacuolar invertase *(Vinv)* and invertase inhibitors in CIS regulation. Transcript levels of *Vinv* **(a)** and invertase inhibitors, INH2ALPHA **(c)** and INVINH1 **(d)** in Bintje, Nico, Laro and Verdi potato tubers after storage at 4° C and 9° C for 1,3 and 5 months, n= 4-5. **b)** VINV protein activity assayed from freeze-dried potato tubers of Bintje, Nico, Laro and Verdi after 3 months of storage at 4° C. n=5 pooled biological replicates. a1, a2, a3 represent statistically significant differences between varieties at P<0.05 computed by Tukey test and * represents significant differences between temperatures at p<0.05 computed by the student’s t-test.

Furthermore, studies have revealed that in addition to the VINV pathway, sucrose is also broken down to UDP-glucose and fructose via the sucrose synthase (SuSy) pathway^13,21^ (Fig. S1A). We investigated the involvement of the SuSy pathway in regulation of CIS-phenotype in CIS contrasting potato varieties Nico and Verdi by RT-qPCR. We observed higher *SuSy* transcript levels in Verdi relative to Nico (Fig. S1B), which inversely correlate with the *Vinv* transcript levels and reducing sugar levels for the respective varieties. These results suggest that the VINV pathway is the principal pathway for breakdown of sucrose to reducing sugars during cold storage.

We subsequently assayed the transcript levels of the two most abundant forms of invertase inhibitors: *INH2ALPHA,* localized predominantly in the vacuole and *INVINH1,* localized mainly in the cytosol^22,25,30^. Generally, we observed higher *INH2ALPHA* transcript levels in Verdi relative to the CIS-susceptible varieties Nico and Laro (Fig. 3C). However, like what was observed for *Vinv,* the transcript level of *INH2ALPHA* in Bintje was as high as that in Verdi. *INVINH1* transcript levels (Fig. 3D) in Verdi was significantly higher than the levels in Bintje and Nico, but similar to the levels in Laro. Combined analysis of the transcript levels of these inhibitors in the potato varieties studied revealed overall higher levels in the CIS-resistant one, Verdi, relative to the CIS-susceptible varieties Bintje, Nico and Laro (Figs. 3C and 3D). Interestingly, the variety accumulating highest levels of reducing sugars, Nico, showed the lowest levels of *INVINH1* and *INH2ALPHA* transcripts. Our results suggest that the concomitant increase in the levels of *INH2ALPHA* and *INVINH1* in Verdi during long term cold storage could contribute to the low level of its VINV activity. All transcript expression data were normalized to the reference gene *18S rRNA,* identified using BestKeeper^31^, to be very stable under these experimental conditions (Fig. S2A and B)

### Genetic components of the *Vinv* promoter regulate CIS

The key role played by the *Vinv* gene in the accumulation of reducing sugars^16^ and the very minimal induction of the *Vinv* gene by cold temperature in Verdi tubers prompted us to investigate regulatory elements of the *Vinv* promoter. A 1720-1730bp region upstream of the ATG-start codon of *Vinv* gene from Bintje, Nico, Laro and Verdi T0 samples (samples collected prior to cold storage) was sequenced and analyzed. Clustal Omega alignment analysis of these sequences revealed high similarity amongst varieties ranging from 98.33 % identity in Bintje vs Laro and Verdi vs Laro, to 99.59 % identity in Verdi vs Bintje (Fig. S3A). However, many SNPs were identified predominantly in the Verdi and Bintje promoters as shown in the alignment of the promoter sequences from all four varieties (Data file S1). Phylogenetic analysis of these sequences revealed that Bintje and Verdi are closely related, with close phylogenetic distances, but distant from Nico and even further distant from Laro (Fig. S3B). Noticeably the phylogenetic relationship matched the pattern of *Vinv* gene expression in the respective varieties (Fig. 3A). Comparative motif analysis of these promoter sequences using PlantCare database^32^ did not reveal any differential motifs associated with the CIS phenotypes in these varieties. However, the SNPs introduced new motifs present solely in Verdi (TATC-Box) or in Verdi and Bintje (Unnamed_6), which could potentially play a role in the regulation of the *Vinv* transcript during cold storage (Fig. S3C). The TATC-Box for example is involved in response to gibberellic acid, a hormone involved in regulation of growth and metabolism during cold stress^33^.

### The 1.0-1.6kb region of *Vinv* promoter is involved in regulation of CIS phenotype in Verdi

*Vinv* promoter fragments of 1.6kb, 1kb and 545bp from each potato variety were used to drive the expression of a luciferase gene cloned in a modified pCambia2300 plasmid (Fig. 4A, Fig. S4). We performed Agrobacterium-mediated transient transformation of potato tuber disks with constructs carrying *Vinv* promoter - luciferase cassettes differing in promoter sequence length. The luminescence intensity after incubation of transformed potato disks at 4° C varied depending on the identity of the *Vinv* promoter. We recorded a positive correlation between promoter length and luciferase activity for promoter sequences originating from Bintje and Laro varieties, indicative of regulatory elements in the distal part of those *Vinv* promoters. A mixed pattern was observed for promoter sequences from Nico and Verdi as luciferase activity peaked with the 1kb promoter sequence and decreased with the 1.6 kb promoter sequences (Fig. 4B). Importantly, the luminescence intensity generated by the luciferase expression under the control of the 1.6 kb *Vinv* promoter sequence from Verdi was lower (averagely 30.35 luminescence units) compared to the 1.6kb promoter sequences from the CIS-susceptible varieties Bintje, Nico and Laro (100-150 luminescence units) (Fig. 4B). This observation suggests that the transient promoter activity assay in potato tuber disks can be used as a proxy to estimate the transcriptional activities of the endogenous potato *Vinv* promoters. Though no specific region could be unambiguously identified in all varieties to be responsible for the regulation of *Vinv* gene expression during cold storage, truncation of the Verdi promoter from 1.6kb to 1kb led to an over 2-fold increase in the luciferase activity (Fig 4B), suggesting that the 1kb-1.6kb region of the promoter is crucial in determining the CIS-phenotype of this variety. In order to investigate variety-dependent regulation of the promoters, we transiently transformed tuber disks from Verdi and Nico varieties with the 1.6kb *Vinv* promoter - luciferase constructs. The 1.6kb *Vinv* promoter sequence from Verdi retained a low transcriptional activity when used to transiently transform Nico tuber disks. Similarly, the 1.6kb fragment of the *Vinv* promoter from Nico retained high transcriptional activity when used to transiently transform Verdi tuber disks (Fig. 4C). These results suggest that the observed low transcriptional activity and the concomitant low VINV activity in Verdi relative to the CIS-susceptible varieties is potentially a result of the inherent genetic makeup of the *Vinv* promoter.

**Fig. 4:**
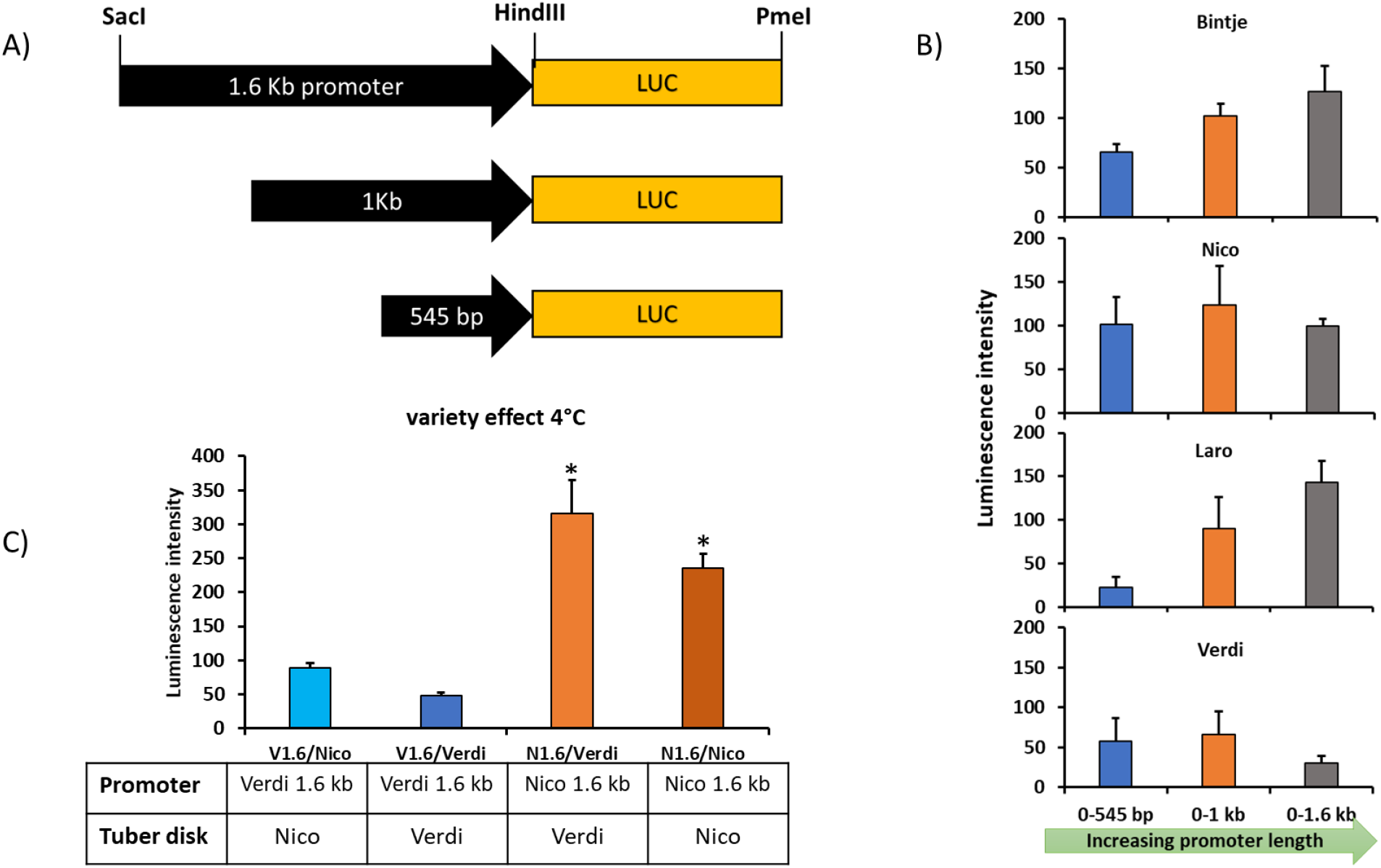
Identification of cold responsive region of *Vinv* promoter. Vinv Promoter-Luciferase truncation constructs (**A**) and luciferase luminescence intensities (**B**) of the different Vinv promoter fragments from Bintje, Nico, Laro and Verdi after storage of transformed tuber discs at 4° C. (**C**) Luciferase assay of *Vinv* promoter activity from CIS-resistant Verdi variety in contrasting CIS-susceptible Nicola potato tubers (V1.6/Nico) and vice versa (N1.6/Verdi). The promoter activities of each promoter in its corresponding tuber (V1.6/Verdi) and (N1.6/Nico) served as respective controls. n=3 and * represents significant differences between test samples and between control samples at P<0.05 using the student’s t-test.

### Differential DNA methylation of the *Vinv* promoter regulates CIS

DNA-cytosine hyper-methylation is known to cause repression of gene expression in plants and mammals^34,35^. DNA methylation was assayed on the promoter region of each potato variety stored for 3 months at 4° C by bisulfite sequencing. Conversion of 91 % of all cytosines to thymines of the internal control *ATP1.1* gene^36^ indicated high and similar efficiency of bisulfite conversion in all samples (Data file S2). In our analysis, Verdi *Vinv* promoter showed higher levels of DNA-cytosine methylation (8.4% non-methylated cytosines) compared with the promoters from Bintje, Nico and Laro (12.9%, 10.3% and 12.3% non-methylated cytosines respectively) (Fig. 5A and Data file S3). Noticeably, Nico showed the highest levels of cytosine methylation amongst the CIS-susceptible varieties. Detailed comparative analysis of the methylation pattern of the *Vinv* promoter revealed that the segment between 1.0-1.7kb showed distinct hypermethylation in Verdi (0.4% non-methylated cytosines) and lowest level of methylation in Nico (9.9% non-methylated cytosines, Fig. 5B). This segment of the *Vinv* promoter is the same segment which showed decreased luciferase activity for Verdi in the truncation experiment (Fig. 4B), thus suggesting that DNA-cytosine methylation, especially in the 1.0-1.7kb region of the Verdi *Vinv* promoter is implicated in the regulation of *Vinv* gene expression under cold conditions, and contributes to the discrepant CIS-phenotypes observed in the potato varieties analyzed in the present study.

**Fig. 5:**
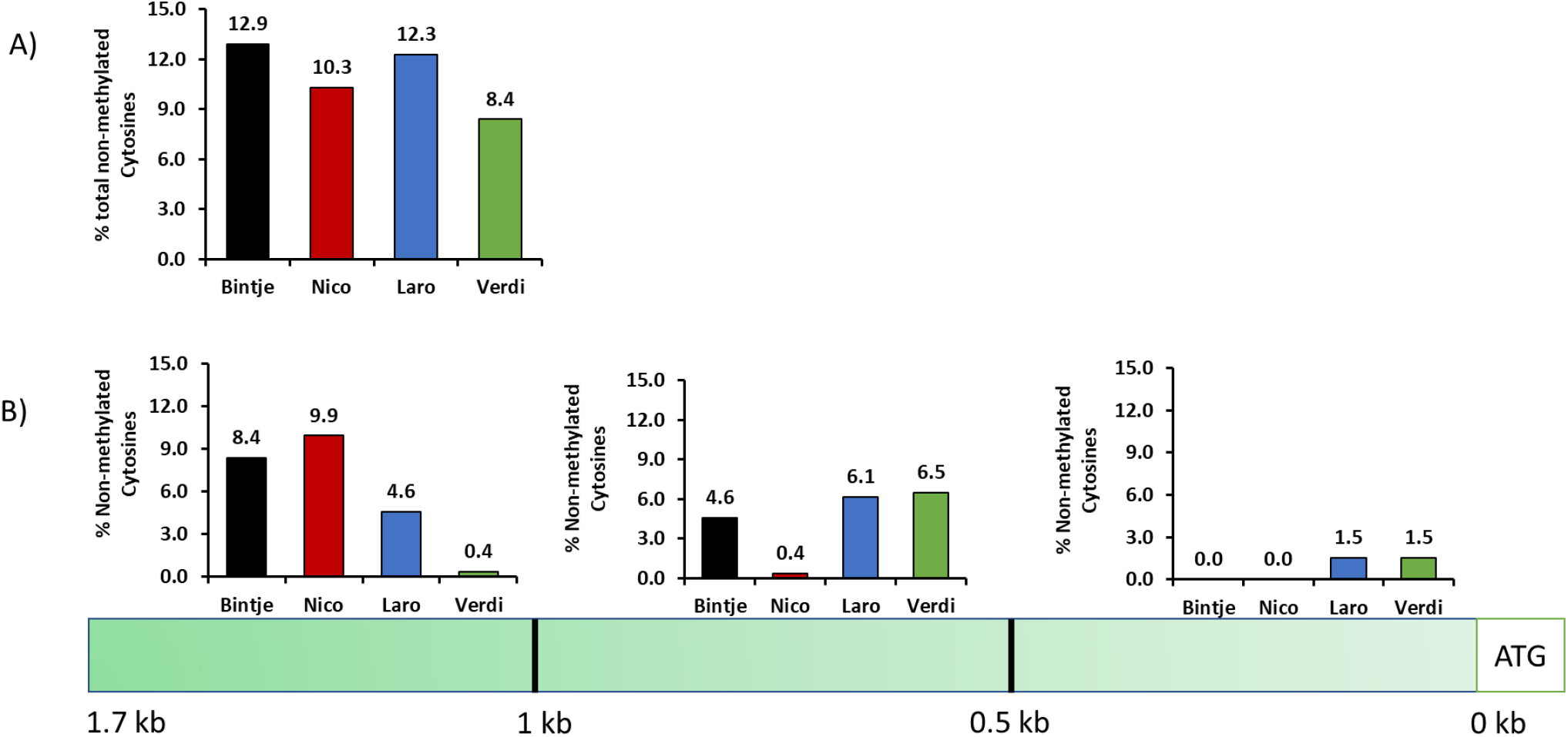
*Vinv* promoter DNA methylation in CIS-phenotype determination. Total (**A**) and segmental (**B**) percentage of non-methylated cytosines on 1.7 kb region of the Vinv promoter from Bintje, Nico, Laro and Verdi potato varieties after bisulfite conversion and sequencing of DNA extracted from tubers stored for 3 months at 4 ° C. The level of non-methylated cytosines of each promoter sequence is represented as the percentage of cytosines converted to thymines (C-T), relative to the total cytosines in the non-bisulfite-treated promoter.

### Evidence of regulation of *Vinv* expression by DNA methylation in the 1.0-1.7kb promoter region

In order to assess the role of DNA hyper-methylation in the 1.0-1.7 kb region of *Vinv* promoter on promoter activity, we performed targeted methylation experiments. The variety Nico was used because it harbors low levels of DNA methylation in the 1.0-1.7 kb target region. We used the CRISPR-dCas9-DRM2 technology^37,38^ to investigate the effect of *de novo* DNA methylation in the 0. 6-1kb and 1.0-1.7kb regions of the *Vinv* promoter on its transcriptional activity. Targeting *de novo* DNA methylation at specific regions of the promoter (Fig. 6A) resulted in lower *Vinv* transcript level relative to their corresponding controls (Fig. 6B). Targeting *de novo* DNA methylation in the 1. 0-1.7 kb promoter region (pC2300-35s-dCas9-AtDRM2-sgRNA2) led to a significant 3-fold decrease in *Vinv* gene expression relative to its control (pC2300-35s-dCas9-sgRNA2). However, this decrease was less remarkable when *de novo* DNA methylation was targeted to the 0.6-1kb promoter region (pC2300-35s-dCas9-AtDRM2-sgRNA3), showing 1.5-folds decrease in *Vinv* gene expression relative to its corresponding control (pC2300-35s-dCas9-sgRNA3) (Fig. 6C): The significant 3-fold decrease in *Vinv* gene expression observed in Nico when DNA methylation was targeted to the 1.0-1.7kb promoter region could be attributed to the availability of more non-methylated cytosines for de-novo DNA methylation (9.9% non-methylated cytosines) compared with the adjacent 0.6-1 kb region (0.4% non-methylated Cytosines). In addition, a possible spread of *de novo* DNA methylation to the 1.0-1.7 region could also account for the slightly decreased *Vinv* expression when targeting the 0.6-1 kb promoter region..

**Fig. 6:**
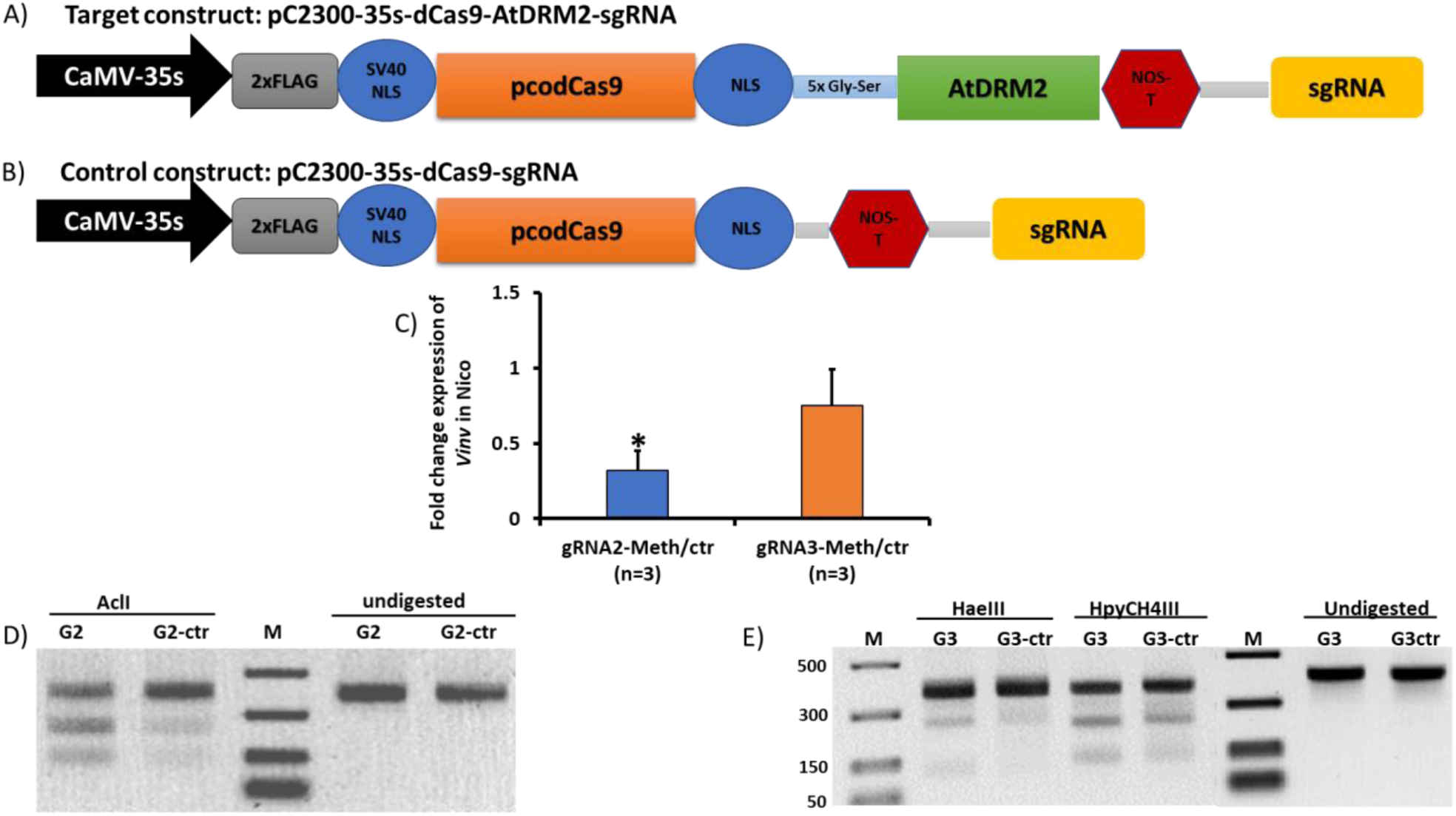
de novo DNA hypermethylation of the *Vinv* promoter. Schematic representation of de novo DNA methylation target construct (A) and control (B). Folds change of *Vinv* expression in de novo methylated tuber disks of Nico variety, relative to controls (non-de novo methylated) (C). Digestion profile of bisulfite PCR amplicons from gRNA2 (gR2) targets and controls using AclI restriction endonuclease (D) and gRNA3(gR3) and controls using HaeIII and HpyCH4III restriction endonucleases (E). * indicates significant difference between target and control at P<0.05 using the student’s t-test. M represents molecular weight marker, and arrowhead ▶ indicates expected bands after digestion

To ascertain that the observed repression of *Vinv* gene expression was as a result of *de novo* DNA methylation in the target regions, bisulfite converted DNA extracted from a pool of three biological replicates each of Nico target and control sample were used to perform PCR using primers spanning approximately 400-450 bp around the sgRNA target site (**Table. S1**). Restriction digestion of target and control DNA amplicons from bisulfite PCR using AclI for sgRNA2 and HaeIII and HypCH4III for sgRNA3 showed consistently higher band intensities in samples targeted for *de novo* DNA methylation compared with their corresponding control samples (Fig. 6D and 6E). *De novo* DNA methylated cytosines were protected from bisulfite conversion to uracil, thus retained the restriction sites which were otherwise lost in non-methylated-bisulfite converted samples. Hence, intense bands were observed in *de novo* methylated samples after restriction digestion relative to their controls. Taken together, these results demonstrate that our CRISPR-dCas9-DRM2 system was efficient in introducing *de novo* DNA methylation in the promoter region of the *Vinv* gene from Nico.

## Discussion

The health dangers associated with acrylamide which accumulates in processed potato products such as fries and chips have forced the European Commission to set benchmark values for acrylamide in potato chips and French fries to 750 μg/kg and 500 μg/kg respectively (Commission Regulation (EU) 2017/2158). With the recent ban on CIPC use in the EU, the potato industry is prompted to find alternative anti-sprouting agents or methods for long term potato storage, including cold storage to reduce sprouting. However, a majority of the consumer- and industry-preferred commercial potato varieties when stored at low temperatures result in processed products with acrylamide content exceeding the benchmark values^28,39^. Our present quantification of reducing sugars accumulating during storage at low temperature along with previous analysis^28^ was instrumental to classify Verdi as CIS-resistant variety and Bintje, Nico and Laro as CIS-susceptible. These contrasting CIS-phenotypes served as the basis of our quest to decipher the underlying molecular mechanisms leading to CIS-resistance in the Verdi variety. Our findings corroborate previous observations that VINV activity is crucial in determining the amount of reducing sugars in potatoes, which in turn determines the level of acrylamide in processed potato products^16^. We observed a good correlation between the transcript levels of *Vinv* gene and the accumulation of reducing sugars in potato tubers of the selected varieties, with the exception of Bintje variety. However, the activity of VINV from all the varieties studied revealed perfect correlation with their corresponding reducing sugar levels, indicating that the regulation of VINV activity occurs at both the transcriptional and post-translational levels. One such post-translational regulation of VINV activity has been reported by invertase inhibitors^22^. We show that the combined transcript levels of the most abundant invertase inhibitor genes, *INVINH1* and *INH2ALPHA,* inversely correlated with the respective VINV activities, thus suggesting a role of the invertase inhibitors in regulating VINV activity and ultimately the CIS-phenotype in the selected potato varieties. Worthy of note, is the fact that Verdi portrayed distinct patterns of sucrose accumulation over time, compared to the CIS-susceptible varieties. Sucrose levels increased with storage time for Verdi but decreased for the CIS-susceptible varieties. Conversion of sucrose to reducing sugars is known to occur simultaneously through two pathways: the VINV pathway and the sucrose synthase (SuSy) pathway^13,21^. The very low activity of the VINV pathway in Verdi forces sucrose to be broken down through the SuSy pathway to produce UDP-Glucose which is subsequently converted back to sucrose through a series of enzymatic reactions (Fig. S1A). This pool of sucrose in combination with the fraction derived from the breakdown of starch is possibly the source of the increasing sucrose content over storage time in Verdi. This hypothesis was confirmed by higher transcript levels of *SuSy* observed in 3- and 5-month stored Verdi potato tubers compared to the corresponding levels in the CIS-susceptible variety Nico (Fig. S1B).

Importantly, this work reveals DNA methylation in the promoter region of *Vinv* as the first line transcriptional regulator of CIS, with DNA methylation in the 1.0-1.7 kb region of the promoter being the major regulatory region during cold storage. We established the CRISPR-dCas9-Methyltransferase system^38^ in potato tuber disks to target *de novo* DNA methylation and to demonstrate that the 1.0-1.7kb region of the *Vinv* promoter is involved in regulating its transcriptional activity during cold storage. In addition to the DNA methylation-based regulation of the *Vinv* promoter, SNPs observed predominantly in the *Vinv* promoter of Verdi and Bintje might also play a distinct role in the regulation of CIS. However, further characterization of these SNPs and the motifs generated as a result of the SNPs are required to ascertain their involvement in determining the CIS phenotypes of these potato varieties. Comparative methylation analysis and targeted *de novo* methylation experiments (Figs 5 and 6) also suggest that the CIS-resistant phenotype can be implemented in CIS-susceptible genotypes by targeted *de novo* methylation of the *Vinv* promoter. While previous efforts have been focused on knockout or knockdown of *Vinv* gene to mitigate CIS in potato tubers, our findings lay the groundwork for artificial epigenetic-based regulation of *Vinv* gene in potato tubers.

## Methods

### Experimental design

Potato varieties were selected based on their importance, commercial use and reported characteristics. Bintje is used predominantly for French fries production, Nico is used for cooking while Laro and Verdi are used for chips production. Freshly harvested tubers of Bintje, Nicola (Nico), Lady Rosetta (Laro) and Verdi potato varieties were sourced from fields in Belgium and Switzerland. These potatoes were kept at room temperature for two weeks before storage at 4° C and 9° C for a period of 5 months. Sampling was done before storage (T0) and after 1, 3 and 5 months of storage. The inner sections of three independent tubers were pooled together to constitute one biological sample. The sample materials were freeze-dried and ground into fine powder using a mortar and pestle. Unless otherwise specified, all experiments were carried out on these samples. Primers employed in this work are listed in Table S1.

### Sugars content

Sucrose, Glucose and fructose contents were determined using the Sucrose, D-Fructose and D-Glucose kit according to the assay procedure (Megazyme, K-SUFRG 04/17) with some modifications. For each sample, 200 mg of freeze-dried powder were suspended in 1mL distilled water and vortexed thoroughly. The suspensions were centrifuged at 15000 rpm for 10 minutes at 4° C and supernatants were collected without disturbing the pellet. Determination of sucrose, glucose and fructose content was then performed as described in the microplate procedure of the Megazyme K-SURFG assay procedure 04/17 using the Microplate 96 wells F-Bottom (Greiner bio-one, Germany). The path length of the microplate was adjusted to 1 cm by dividing the absorbance with the value of 0.625, which is a function of the diameter of the well and the total reaction volume. Glucose, fructose and sucrose contents were calculated as described in the assay procedure.

### RNA extraction and RT-qPCR

RNA was extracted from freeze-dried tuber samples using the phenol-chloroform protocol as previously described^40^ with some modifications. Approximately 100 mg of starting material was used, and the volumes of reagents (NaCl SDS, Na_2_SO_3_, TBE, and beta-mercaptoethanol) were scaled down 5 folds. RNA quality was verified on 1% agarose gel and intact RNA was treated with DNAseI (Bioke, Netherlands) according to the manufactures’ instructions. Aliquots of DNAse treated RNA were again run on a 1% agarose gel to ensure complete elimination of genomic DNA. First strand cDNA was synthesized from 500 ng of each DNA-free RNA sample using GoTaq Reverse Transcription, Oligo dT kit (Promega, USA) according to the manufacturers’ instructions. cDNA was diluted 5 folds with distilled water and used in a reaction mix composed of 1x GoTaq qRT-PCR master mix (Promega, USA) and 10 nM primers for RT-qPCR. Primer sequences are listed in Table S1. Each sample was analyzed in triplicates on a CFX96 Real-Time System (BioRad, USA) using the following program: Initial denaturation at 95° C for 3 mins, then 40 cycles of 95° C for 10 sec (denaturation), 60° C for 30 sec (Annealing) and 72° C for 30 sec (elongation), followed by a plate read. The PCR cycles were followed by generation of a melting curve between 65° C and 95° C with 0.5° C increments. Relative gene expression was calculated using the comparative cycle threshold method^41^, and expression levels of candidate genes were normalized to the mean delta cycle threshold (dCT) of the housekeeping gene. In order to identify the best stable housekeeping gene(s) for normalization, primers for elongation factor 1-α *(EF1α), 18S rRNA,* adenine phosphoribosyl transferase *(APRT),* cytoplasmic ribosomal protein L2 (L2), *Peroxin* and *Profilin* genes (Table S1) were used to perform RT-qPCR on biological triplicates of Bintje, Nico, Laro and Verdi tuber samples stored for 3 months at 4° C and 9° C. The stability of these housekeeping genes was analyzed with the help of the BestKeeper software^31^.

### Acrylamide content

Laro and Verdi tubers were grown and stored at 4° C and controlled CO_2_ conditions for 5 months in Switzerland, after which ten tubers of each variety were transformed to chips by Agroscope-Switzerland. Just before processing to chips, the glucose content of three tubers from each variety was measured using an Accu-Check blood glucometer. The chips colors from Verdi tubers were visually compared to those of Laro tubers. The chips obtained from the three biological replicates of each variety were manually ground to powder in a sealed plastic bag using a rolling pin. Acrylamide was extracted from each sample in duplicates, and quantified by GC-MS on a Trace GC/Trace DSQ instrument (Thermo Scientific, Dreieich, Germany) in chemical ionization mode exactly as described previously^43^.

### Vacuolar invertase activity

Vacuolar invertase activity was assessed using approximately 300 mg of freeze-dried samples from potatoes stored for 3 months at 4° C. Approximately 60 mg each of five biological replicates for each variety were pooled to form a single sample. Protein extraction, desalting and enzyme assay was performed exactly as described by Bhaska and co-workers^16^. The glucose content of samples and controls was quantified using the K-SUFRG kit as described in the assay procedure (Megazyme, K-SUFRG 04/17), and the amount of glucose produced was determined by calculating the difference in glucose content between samples and controls. Total protein content for each sample was quantified using the Bradford method^44^ and vacuolar invertase activity was calculated as glucose concentration produced per hour per microgram of total protein.

### Asparagine content

Free Asparagine content was quantified in 3 months stored potato samples with the help of the L-Asparagine/L-Glutamine/Ammonia kit as described in the assay procedure (Megazyme, K-ASNAM 07/17), with some modifications. Approximately 200 mg of each freeze-dried sample (50 folds less) was resuspended in 1.6 mL of 1M perchloric acid. After thorough mixing and centrifugation at 15000 rpm for 10 minutes at 4° C, 1 mL of the supernatant was harvested into a 2 mL tube and pH was adjusted to 8.0 with 2 M KOH. The volume was then adjusted to 1.5 mL with distilled water. The subsequent procedures of quantification by the microplate format and calculations of asparagine content were performed as described in the assay procedure K-ASNAM 07/17, with adjustment of the path length to 1 cm.

### Illumina and Bisulfite sequencing

High quality genomic DNA was extracted from freeze-dried tubers using the CTAB-chloroform-phenol extraction method. Approximately 30 mg each of 3 biological replicates of each sample type (variety/temperature) were pooled to form a single sample for DNA extraction. Bisulfite conversion was performed using the methyl edge bisulfite conversion system exactly as described by the manufacturer (Promega, USA). Four overlapping PCR primers (Bs-1F/1R, Bs-2F/2R, Bs-3F/3R, Bs-4F/4R, Table S1) were designed to cover approximately 1.7 kb of the *Vinv* promoter region upstream the ATG start codon. These primers were designed avoiding C-G rich regions and illumina adapters were added to the 5’-ends of each forward and reverse primer. Bisulfite PCRs were performed on the bisulfite converted samples using zymo-Taq polymerase (BaseClear, Netherlands), and gel purified fragments from individual samples were pooled together for illumina sequencing. T0 samples were not subjected to bisulfite conversion and served as control for the converted samples. Five overlapping PCR primer pairs (Vinvpro_1980-1540-F/R, Vinvpro_1574-1211-F/R, Vinvpro_1287-987-F/R, Vinvpro_1007-524-F/R and Vinvpro_547-66-F/R, **Table S1**) were designed to cover approximately 1.7 kb region of the *Vinv* promoter, with the respective illumina adapters added to their 5’-ends. PCR was performed using Q5 DNA polymerase (NEB). A 227 bp fragment of the mitochondrial ATPase subunit-1 gene was amplified from the bisulfite converted and non-converted DNA samples using the degenerate ATP1.1-F and ATP1.1-R primer pair described previously^36^. This fragment served as internal control for bisulfite conversion. Illumina sequencing was performed on both bisulfite converted and non-converted PCR products using the 2*300 bp paired end MiSeq (GIGA platform, University of Liège, Belgium). Approximately 300,000 reads were generated per sample. Reads were assembled and analyzed using CLC-Bio software (Qiagen, USA). Sequences from the different potato varieties were aligned with the help of the pairwise alignment tool for CLC-Bio and the Clustal Omega sequence alignment tool and genetic variation among the different potato varieties were identified by comparing T0 unconverted samples. Comparative motif analysis was performed using the PlantCARE software^32^. Bisulfite converted sequences were aligned with each other and with their corresponding non-converted sequences, and the methylation percentage was calculated.

### Generation of vectors for transformation

#### *Vinv* promoter-LUC construct generation

Promoter-Reporter constructs were generated for different lengths of the *Vinv* promoter, with luciferase as the reporter gene. Three promoter length; 545 bp, 1 kb and 1.6 kb upstream the ATG start codon were amplified from genomic DNA of each variety by PCR, using the primer pairs *StVinv*-promoter_545-F/Common-R, *StVinv*-promoter_1kb-F/Common-R and *StVinv*-promoter_1.6kb-F/common-R, respectively. SacI and HindIII restriction site sequences were added at the 5’-ends of the forward and reverse primers, respectively (Table S1). The firefly luciferase sequence was amplified from the PsiCHECK2 vector (Promega), using LUC-F and LUC-R primers containing respectively HindIII and PmeI flanking restriction sites at their 5’-ends. The gel-purified promoter and luciferase fragments were cloned independently in to the Pjet1.2 blunt vector (Fisher Scientific). Positive colonies were identified by colony PCR and confirmed by sanger sequencing using PJET-F primer. The promoter fragments were then digested out of the respective Pjet1.2-recombinant plasmids by use of SacI and HindIII restriction enzymes and cloned into the pCambia2300 vector to generate the plasmids pC2300-Promoter. The Luciferase fragment was subsequently digested from the PJet1.2-recombinant plasmid with the help of HindIII and PmeI restriction enzymes and cloned in to the pC2300-promoter plasmid to generate the PC2300-Promoter-LUC plasmid. Finally, the CaMV-35S promoter driving the Kanamycin resistance gene in the pC2300-Promoter-LUC plasmid was removed by digestion with NcoI and Eco53kI restriction enzymes and replaced with a 356 bp fragment from Pjet1.2 obtained by digesting Pjet1.2 with NcoI and PmeI restriction enzymes. These complete constructs (pC2300-Promoter-LUC*) were verified by digestion and sequencing, and then used to transform *Agrobacterium tumefaciens* LBA4404 strain.

#### *De novo* DNA methylation constructs

Guide RNAs (gRNA) were designed in the promoter regions 1.0-1.7 kb (gRNA2) and 0.6-1kb (gRNA3) using the CRISPR RGEN Cas-Designer Tool^45^ and off targets were double checked by blast analysis against the potato genome on phytozome. Assembly of the gRNA cassette (AtU6promoter-Target-gRNA scaffold-Terminator) was performed by PCR using the primer pairs BspEI-gRNA2-F/BspEI-Common-R (for gRNA2) and BspEI-gRNA3-F/BspEI-Common-R (for gRNA3) and a synthesized gRNA cassette as template. Each gRNA was cloned in to Pjet1.2 blunt plasmid to form pJet-gRNA2 and pJet-gRNA3 plasmids, respectively.

CaMV-35s promoter was amplified from pCambia2300 plasmid using the primer pair ApaI-EcoRI-35s-F and NcoI-35s-R and cloned in the ApaI/NcoI sites of pYPQ152 plasmid harboring the plant codon optimized deactivated Cas9 (Pco-dCas9) coding sequence (Addgene Plasmid #69303) to generate the pYPQ152_35s plasmid. The coding sequence of *Arabidopsis thaliana* domain rearranged methyltransferase II (AtDRM2) was amplified from *A. thaliana* Columbia-0 (Col-0) cDNA using the primer pair SalI-DRM2-F and AatII-DRM2-R and cloned into the SalI/AatII site of the pYPQ152_35s plasmid to generate the pYPQ152_35s-AtDRM2 plasmid. The pYPQ152_35s-AtDRM2 plasmid was subsequently digested with EcoRI, and the 35s-dCas9-DRM2 fragment was recovered and cloned into the EcoRI site of pCambia2300 to form pC2300-35s-dCas9-DRM2 plasmid.

The final constructs pC2300-35s-dCas9-DRM2-gRNA2 and pC2300-35s-dCas9-DRM2-gRNA3 were generated by cloning each gRNA fragment from its respective pJet-gRNA plasmid into the BspEI site of pC2300-35s-dCas9-DRM2 plasmid. Control plasmids were constructed similarly to the final constructs but lacking the methyltransferase DRM2 sequence. The 35s-dCas9 fragment from pYPQ152_35s plasmid was extracted by EcoRI/SalI digestion and cloned in to pC2300 to produce pC2300-35s-dCas9. Undigested PCR purified gRNA fragments were cloned in to pC2300-35s-dCas9 at the PmeI site to produce respectively pC2300-35s-dCas9-gRNA2 and pC2300-35s-dCas9-gRNA3 control plasmids.

### Agrobacterium-mediated transformation of potato disks

Agrobacterium LBA4404 harboring the *Vinv* promoter-LUC constructs: pC2300-1.6kb-LUC*, pC2300-1.0kb-LUC*, pC2300-545bp-LUC* from Bintje, Nico, Laro and Verdi varieties and the de novo DNA methylation constructs: pC2300-35s-dCas9-DRM2-gRNA2, pC2300-35s-dCas9-DRM2-gRNA3, pC2300-35s-dCas9-gRNA2 and pC2300-35s-dCas9-gRNA3 were used to transform thin sliced potato tuber disks. Tubers transformed with agrobacterium LBA4404 not carrying a binary vector served as control for the *Vinv* promoter-LUC transformations. A cork borer was used to bore cylindrical portions from potato tubers and a razor blade was used to cut the cylinders in to thin disks of approximately similar thickness. Agrobacteria carrying the constructs were grown to OD600 of 0.9 in LB-Rifampicin (20 mg/L) medium. After centrifugation, the pellets were resuspended in 10 ml of Ms-liquid medium supplemented with acetosyringone to a final concentration of 100 μM. The potato disks were transformed by incubation with the Agrobacterium suspension for 20 minutes, after which the disks were allowed to dry in a sterile laminar hood on sterile filter paper, before transferred to MS-solid medium for co-cultivation at 28° C. After 2 days of co-cultivation, excess Agrobacterium was washed twice for 3 minutes with sterile distilled water, then once for 5 minutes with sterile distilled water containing 300 mg/L cefotaxime. The potato disks were again dried as described above and placed on new MS-solid media layered with sterile surgical mesh on the surface. Each biological sample (ten potato disks from a single tuber constituted a biological replicate) were incubated at 4° C. After seven days of incubation, the tuber disks were rapidly dried with absorbent paper and ground in a mortar under liquid nitrogen. Approximately 300 mg of each ground sample was rapidly weighed in to pre-chilled 2 ml tubes to avoid thawing.

### Luciferase assay of promoter activity

Potato tuber disks (Var. Lucilla) transformed with agrobacterium harboring the *Vinv* promoter-LUC constructs were harvested as described above. One mL of 1X cell lysis buffer from the luciferase assay system (Promega) was added to each sample, thoroughly vortexed to mix and gently agitated on a shaker for 2 hours at 4° C. The suspension was then centrifuged at 15000 rpm for 10 minutes at 4° C and the supernatant were collected in new 1.5 ml tubes placed on ice. The luciferase assay was performed by thoroughly mixing 10 μl of a sample with 50 μl of the luciferin substrate (Luciferase Assay System, Promega) and measuring the luminescence intensity after 10 and 20 seconds using a luminometer (TD-20/20 Luminometer, Turner Designs, USA). Similar procedures were performed to assess the promoter activity of pC2300-N1.6kb-LUC* in Verdi tuber disks, pC2300-V1.6kb-LUC* in Nico tuber disks, pC2300-N1.6kb-LUC in Nico tuber disks and pC2300-V1.6kb-LUC* in Verdi tuber disks

### *De novo* DNA methylation using CRISPR-dCas9-Methyltransferase system

LBA4404 agrobacterium harboring the respective plasmids for targeted DNA methylation and controls were used to transform thin potato tuber disks of Nico and Verdi. From each biological replicate, samples for DNA and RNA extraction were harvested. RNA was extracted and cDNA was synthesized from the harvested samples. *Vinv* gene expression was assayed in target samples and controls by RT-qPCR using the *StVinv-F* and -R primers and *dCas9-F* and -R primes (Table S1) as housekeeping gene. Upon transformation of potato tuber disks with agrobacteria carrying the target constructs and controls, only the cells on the surface having contact with the agrobacterium have the potential to be transformed. Very thin slices of the tuber disks were used to minimize the amount of non-transformed cells. In addition, *Vinv* gene expressions in these experiments were normalized to the corresponding *dCas9* transcript as housekeeping gene, to eliminate the bias introduced by non-transformed cells.

DNA was extracted from a pool of the three biological replicates of each Nico sample and bisulfite conversion was performed. PCR was performed on the bisulfite converted samples targeted for *de novo* DNA methylation at the 1.0-1.7 kb region (gRNA2) and corresponding controls using Bs-2.1New-F and -R primers (Table S1). Bisulfite converted samples targeted for *de novo* DNA methylation at the 0.6-1kb region (gRNA3) and corresponding controls were amplified using Bs-4-F and -R primers (Table S1). The level of DNA methylation between *de novo* methylation targeted (gRNA2) and corresponding control was assessed by digestion of 50 ng of each bisulfite PCR product with AclI (gRNA2) and HaeIII and HpyCH4III (gRNA3) restriction endonucleases for 2 hours.

### Statistical analysis

The students t-test and the Tukey test were employed for computation of statistically significant difference between varieties and between storage temperatures for data generated in this work.

## Acknowledgements

We wish to thank Dr. Klaus Vosmann (Max Rubner-Institut) for excellent support with GC-MS detection of acrylamide and Lu Hejun (University of Liège) for technical assistance to generate the constructs.

## Author contributions

**LS:** Conceptualization, Methodology, Investigation, Writing-original draft, Reviewing and Editing

**MV:** Investigation, Reviewing and Editing

**ES** and **IS**: Investigation, Review and Editing

**BD:** Reviewing and Editing, Funding acquisition

**HV:** Conceptualization, Methodology, Writing-review and Editing, Supervision, Funding acquisition

## Supplementary Figures

**Fig. S1:**
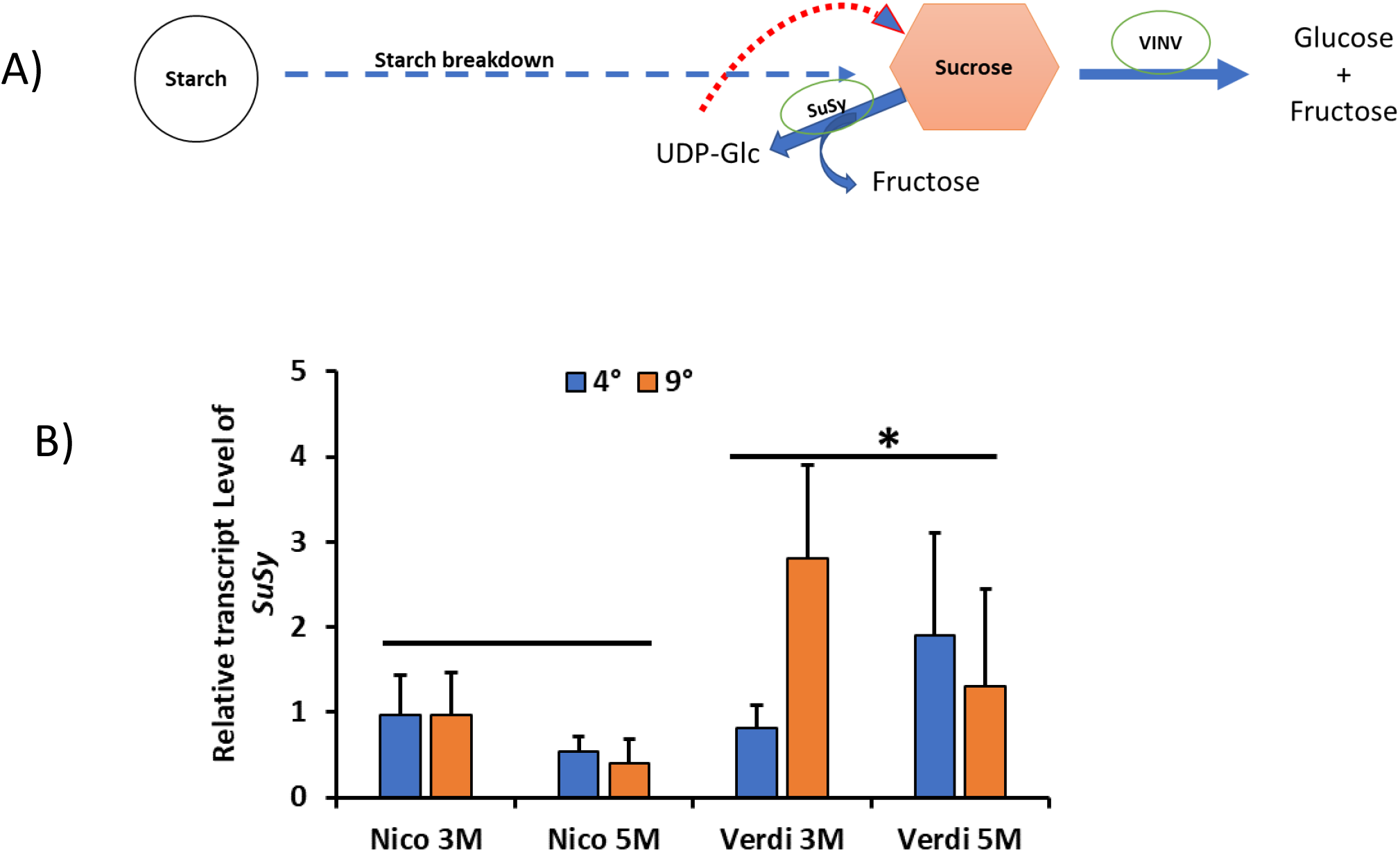
SuSy pathway in CIS. Simplified scheme of starch and sucrose metabolism (**A**) showing the major enzymes involved in the breakdown of sucrose (SuSy and VINV), and (**B**) relative expression of SuSy in contrasting CIS varieties, Nico and Verdi after 3 and 5 months of storage (3M, 5 M). n=5. * indicates significant difference between Nico and Verdi at P<0.05 using the students t-test.

**Fig. S2:**
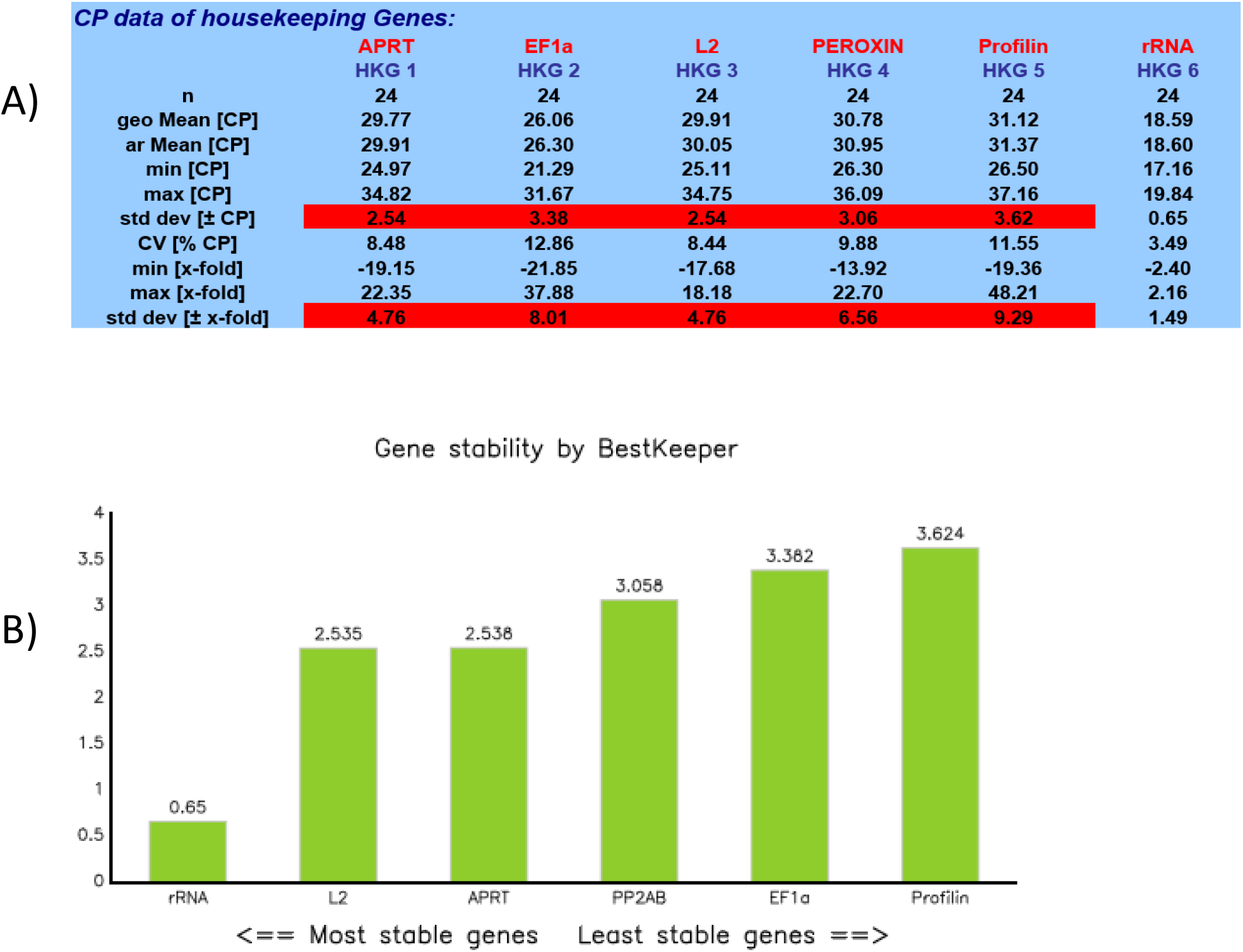
Identification of most stable housekeeping gene(s). BestKeeper analysis of housekeeping gene for stability (A) and ranking of these genes in order of stability (B).

**Fig. S3:**
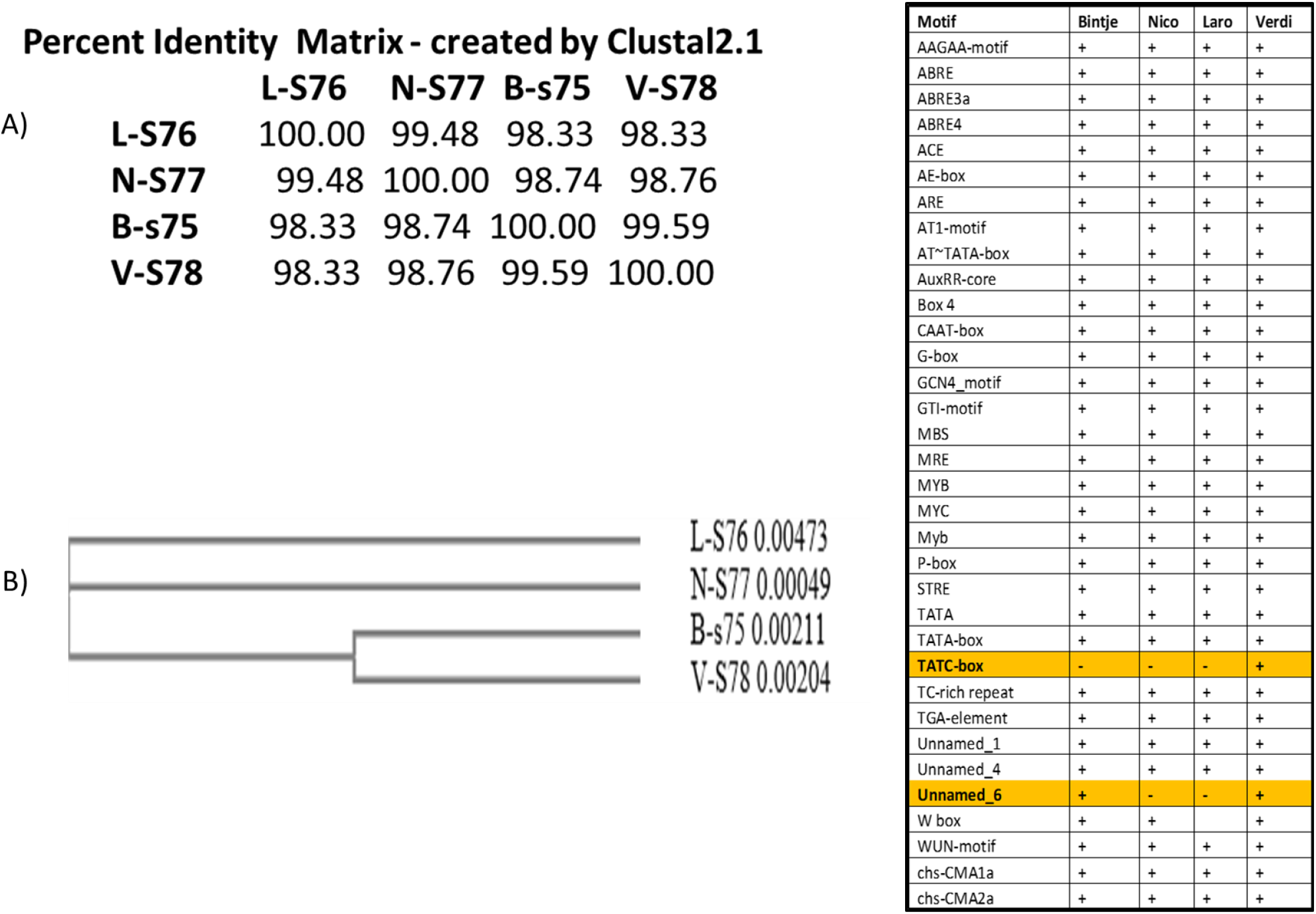
*Vinv* promoter sequence analysis. Comparative analysis of 1.7 kb concensus sequences of promoter regions upstream the start codons of *Vinv* gene from Bintje (B-s75), Nico (N-S77) LaroLaro (L-S76) and Verdi (V-S78). Concensus sequences were generated from ~250000 illumine reads per DNA sample extracted from a pool of 5 biological samples. **A)** percentage identity between sequences, **(B)** cladogram representing phylogenetic distances between sequences generated from sequence alignment using Clustal2.1. **C)** Comparative motif analysis using PlantCare. + and - represent presence and absence of motif respectively. Highlighted motifs are differentially present/absent in different varieties.

**Fig. S4:**
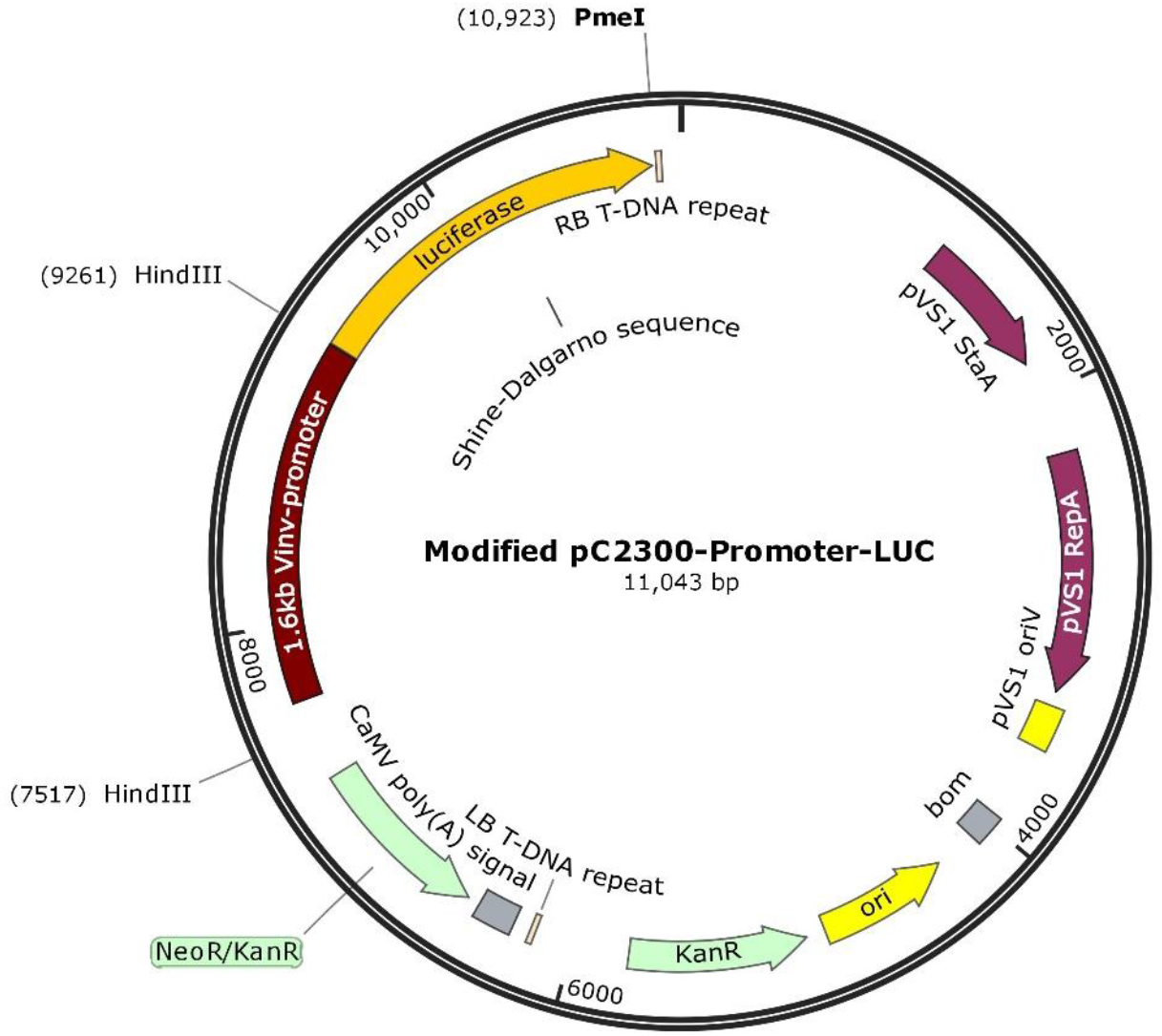
Plasmid map. pC2300-1.6kb-LUC* construct used in the promoter-reporter system, constructed using Snapgene Viewer. The CaMV-35s promoter driving expression of the Kan^R^ gene in the original pC2300 plasmid was replaced by a fragment from pJet1.2.

## Supplementary Table

**Table S1:**
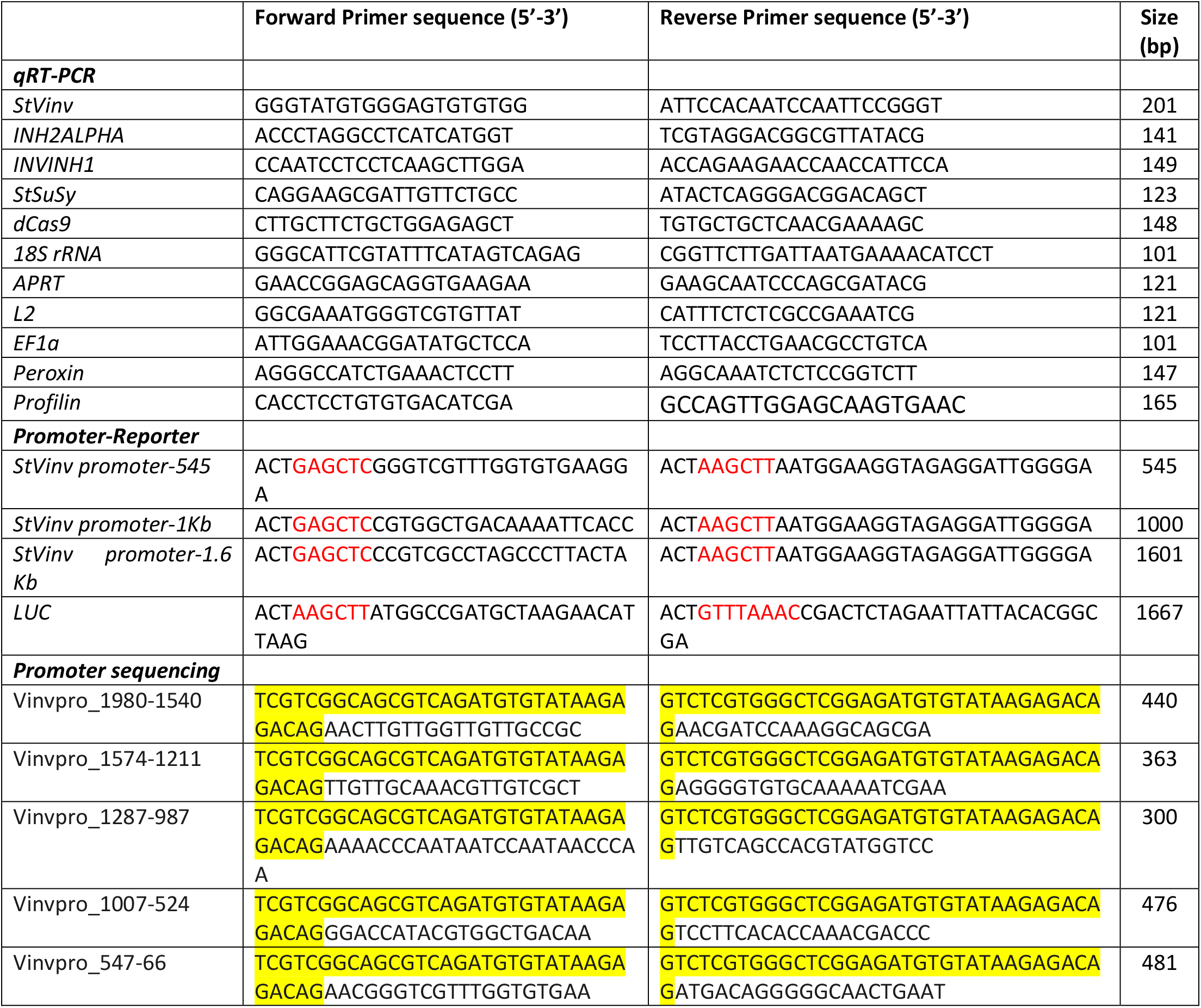
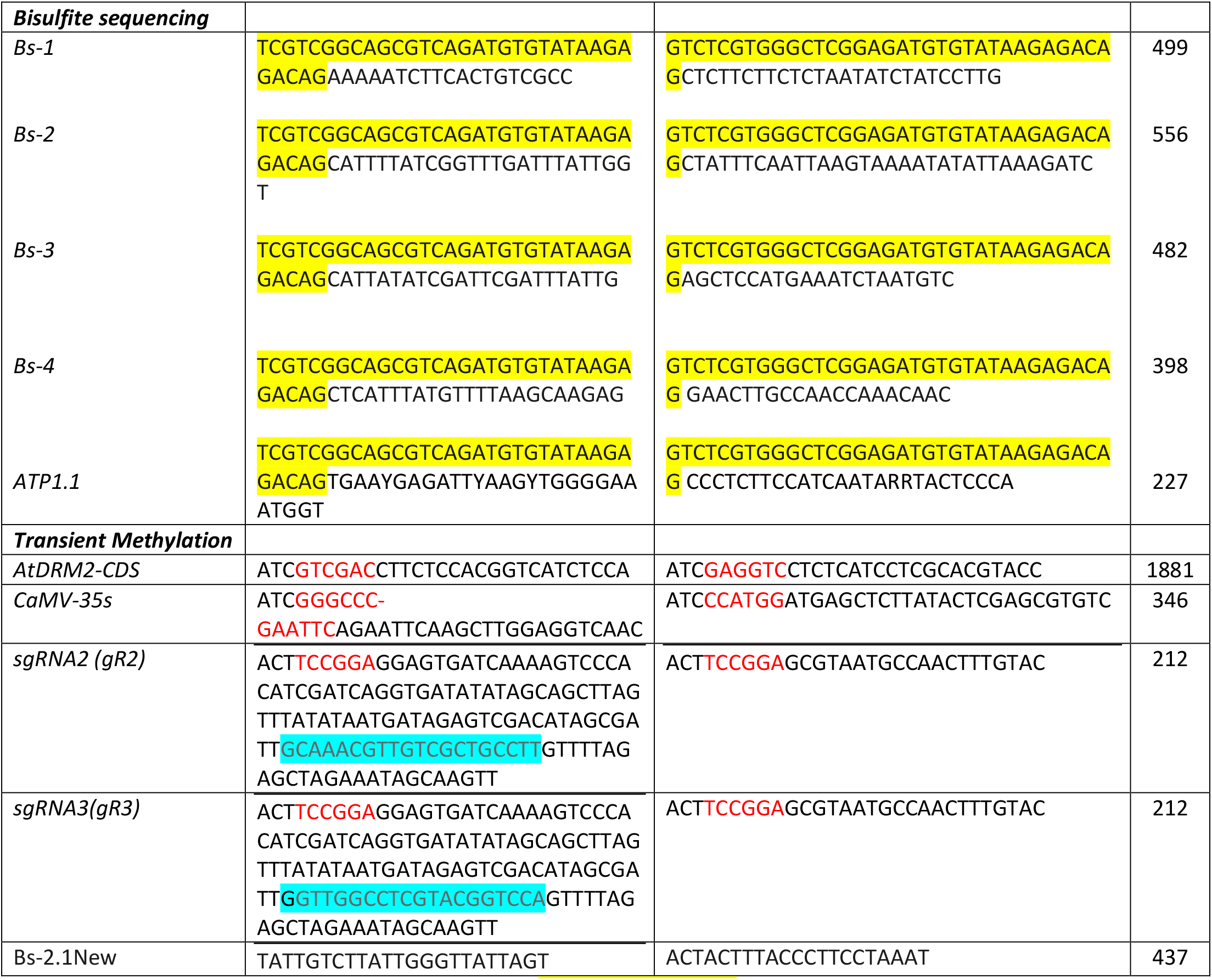
Primer sequences used in this work. Yellow highlighted sequences represent the illumina adapters, Blue highlighted sequences represent the 20-nt sgRNA sequence, and red coloured sequences are sequences of restriction enzymes added to the 5’-end of the primers as overhangs.

## Supplementary data files

**Data file S1**: Alignment ATP1.1-ctr Vs Bs-ATP1.1 Bintje, Nico, Laro,Verdi.

**Data file S2**: *Vinv* promoter alignment from Bintje, Nico, Laro and Verdi T0 samples

**Data file S3:** Data file S3. Merged alignments of unconverted promoter sequences Vs their corresponding bisulfite converted sequences

## References

1. Fujitani, T., Tada, Y. & Yoneyama, M. Chlorpropham-induced splenotoxicity and its recovery in rats. Food Chem. Toxicol. 42, 1469–1477 (2004).

2. Sher Mohammed, N., Flowers, T. & Duncan, H. HPLC-UV Method for the Analysis of Potato Sprout Inhibitor Chlorpropham and Its Metabolite 3-Chloroaniline in Potatoes. 9, 78–85 (2015).

3. Worobey, B. L. & Sun, W.-F. Isolation and identification of chlorpropham and two of its metabolites in potatoes by GC-MS. Chemosphere 16, 1457–1462 (1987).

4. Kleinkopf, G. E., Brandt, T. L., Frazier, M. J. & Möller, G. CIPC residues on stored Russet Burbank potatoes: 1. Maximum label application. Am. Potato J. 74, 107–117 (1997).

5. Daniels-Lake, B. J. The Combined Effect of CO2 and Ethylene Sprout Inhibitor on the Fry Colour of Stored Potatoes (Solanum tuberosum L.). Potato Res. 56, 115–126 (2013).

6. Richard Knowles, N., Knowles, L. O. & Haines, M. M. 1,4-Dimethylnaphthalene treatment of seed potatoes affects tuber size distribution. Am. J. Potato Res. 82, 179–190 (2005).

7. Knowles, L. O. & Knowles, N. R. Toxicity and Metabolism of Exogenous α,β-Unsaturated Carbonyls in Potato (Solanum tuberosum L.) Tubers. J. Agric. Food Chem. 60, 11173–11181 (2012).

8. De Blauwer, V., Demeulemeester, K., Demeyere, A. & Hofmans, E. Maleic hydrazide: sprout suppression of potatoes in the field. Commun. Agric. Appl. Biol. Sci. 77, 343–351 (2012).

9. Coleman, W. K., Lonergan, G. & Silk, P. Potato Sprout Growth Suppression by Menthone and Neomenthol, Volatile Oil Components ofMinthostachys, Satureja, Bystropogon, andMentha Species. Am. J. Potato Res. 78, 345–354 (2001).

10. Şanli, A. & Karadoğan, T. Carvone Containing Essential Oils as Sprout Suppressants in Potato (Solanum tuberosum L.) Tubers at Different Storage Temperatures. Potato Res. (2019). doi:10.1007/s11540-019-9415-6

11. Finger, F. et al. Action of Essential Oils on Sprouting of Non-Dormant Potato Tubers. Brazilian Arch. Biol. Technol. 61, (2018).

12. Dolničar, P., Mavric plesko, I. & Meglic, V. Long-Term Cold Storage Suppress the Development of Tuber Necrosis Caused by PVYNTN. Am. J. Potato Res. 88, (2011).

13. Sowokinos, J. R. Biochemical and molecular control of cold-induced sweetening in potatoes. Am. J. Potato Res. 78, 221–236 (2001).

14. Mottram, D. S., Wedzicha, B. L. & Dodson, A. T. Acrylamide is formed in the Maillard reaction. Nature 419, 448–449 (2002).

15. Pedreschi, F., Moyano, P., Kaack, K. & Granby, K. Color changes and acrylamide formation in fried potato chips. Food Res. Int. 38, 1–9 (2005).

16. Bhaskar, P. B. et al. Suppression of the vacuolar invertase gene prevents cold-induced sweetening in potato. Plant Physiol. 154, 939–948 (2010).

17. Zhang, H. et al. The potato amylase inhibitor gene SbAI regulates cold-induced sweetening in potato tubers by modulating amylase activity. Plant Biotechnol. J. 12, 984–993 (2014).

18. P. Spychalla, J., E. Scheffler, B., Sowokinos, J. & W. Bevan, M. Cloning, Antisense RNA Inhibition, and the Coordinated Expression of UDP-Glucose Pyrophosphorylase with Starch Biosynthetic Genes in Potato Tubers. J. Plant Physiol. 144, 444–453 (1994).

19. Rommens, C. M., Ye, J., Richael, C. & Swords, K. Improving Potato Storage and Processing Characteristics through All-Native DNA Transformation. J. Agric. Food Chem. 54, 9882–9887 (2006).

20. Hou, J. et al. Amylases StAmy23, StBAM1 and StBAM9 regulate cold-induced sweetening of potato tubers in distinct ways. J. Exp. Bot. 68, 2317–2331 (2017).

21. Zhang, H. et al. The roles of starch metabolic pathways in the cold-induced sweetening process in potatoes. Starch - Stärke 69, 1600194 (2017).

22. Mckenzie, M. J., Chen, R. K. Y., Harris, J. C., Ashworth, M. J. & Brummell, D. A. Post-translational regulation of acid invertase activity by vacuolar invertase inhibitor affects resistance to cold-induced sweetening of potato tubers. Plant. Cell Environ. 36, 176–185 (2013).

23. Ou, Y. et al. Promoter regions of potato vacuolar invertase gene in response to sugars and hormones. Plant Physiol. Biochem. 69C, 9–16 (2013).

24. Lin, Y. et al. Interaction proteins of invertase and invertase inhibitor in cold-stored potato tubers suggested a protein complex underlying post-translational regulation of invertase. Plant Physiol. Biochem. 73, 237–244 (2013).

25. Brummell, D. A. et al. Induction of vacuolar invertase inhibitor mRNA in potato tubers contributes to cold-induced sweetening resistance and includes spliced hybrid mRNA variants. J. Exp. Bot. 62, 3519–3534 (2011).

26. Groza, H. I. et al. White Pearl—A chipping potato variety with high level of resistance to cold sweetening. Am. J. Potato Res. 83, 259–267 (2006).

27. Ali, A., Iqbal, M., Ali, Q., Razzaq, A. & Nasir, I. A. Gene profiling for invertase activity: assessment of potato varieties for resistance towards cold induced sweetening. Adv. Life Sci. 3, 63–70 (2016).

28. Elmore, J. S. et al. Acrylamide in potato crisps prepared from 20 UK-grown varieties: Effects of variety and tuber storage time. Food Chem. 182, 1–8 (2015).

29. Chawla, R., Shakya, R. & Rommens, C. M. Tuber-specific silencing of asparagine synthetase-1 reduces the acrylamide-forming potential of potatoes grown in the field without affecting tuber shape and yield. Plant Biotechnol. J. 10, 913–924 (2012).

30. Shivalingamurthy, S. G. et al. Identification and Functional Characterization of Sugarcane Invertase Inhibitor (ShINH1): A Potential Candidate for Reducing Pre- and Post-harvest Loss of Sucrose in Sugarcane. Front. Plant Sci. 9, 598 (2018).

31. Pfaffl, M. W., Tichopad, A., Prgomet, C. & Neuvians, T. P. Determination of stable housekeeping genes, differentially regulated target genes and sample integrity: BestKeeper - Excel-based tool using pair-wise correlations. Biotechnol. Lett. 26, 509–515 (2004).

32. Lescot, M. et al. PlantCARE, a database of plant cis-acting regulatory elements and a portal to tools for in silico analysis of promoter sequences. Nucleic Acids Res. 30, 325–327 (2002).

33. Achard, P. et al. The Cold-Inducible CBF1 Factor–Dependent Signaling Pathway Modulates the Accumulation of the Growth-Repressing DELLA Proteins via Its Effect on Gibberellin Metabolism. Plant Cell 20, 2117 LP – 2129 (2008).

34. Berdasco, M. et al. Promoter DNA hypermethylation and gene repression in undifferentiated Arabidopsis cells. PLoS One 3, e3306–e3306 (2008).

35. Baylin, S. B. & Herman, J. G. DNA hypermethylation in tumorigenesis: epigenetics joins genetics. Trends Genet. 16, 168–174 (2000).

36. Wang, J. et al. Universal endogenous gene controls for bisulphite conversion in analysis of plant DNA methylation. Plant Methods 7, 39 (2011).

37. Vojta, A. et al. Repurposing the CRISPR-Cas9 system for targeted DNA methylation. Nucleic Acids Res. 44, 5615–5628 (2016).

38. Papikian, A., Liu, W., Gallego-Bartolomé, J. & Jacobsen, S. E. Site-specific manipulation of Arabidopsis loci using CRISPR-Cas9 SunTag systems. Nat. Commun. 10, 729 (2019).

39. Muttucumaru, N. et al. Acrylamide-forming potential of potatoes grown at different locations, and the ratio of free asparagine to reducing sugars at which free asparagine becomes a limiting factor for acrylamide formation. Food Chem. 220, 76–86 (2017).

40. Kumar, G. N. M., Iyer, S. & Knowles, N. R. Extraction of RNA from Fresh, Frozen, and Lyophilized Tuber and Root Tissues. J. Agric. Food Chem. 55, 1674–1678 (2007).

41. Livak, K. J. & Schmittgen, T. D. Analysis of Relative Gene Expression Data Using Real-Time Quantitative PCR and the 2-ΔΔCT Method. Methods 25, 402–408 (2001).

42. Misener, G. C., Gerber, W. A., Tai, G. & Embleton, E. J. Measurement of glucose concentrations of potato extract using a blood glucose test strip. Can. Agric. Eng. 38, 59–62 (1996).

43. Haase, N. U., Grothe, K.-H., Matthäus, B., Vosmann, K. & Lindhauer, M. G. Acrylamide formation and antioxidant level in biscuits related to recipe and baking. Food Addit. Contam. Part A 29, 1230–1238 (2012).

44. He, F. Bradford Protein Assay. Bio-protocol 1, e45 (2011).

45. Park, J., Bae, S. & Kim, J.-S. Cas-Designer: a web-based tool for choice of CRISPR-Cas9 target sites. Bioinformatics 31, 4014–4016 (2015).

