## Supplementary material for "Differential DNA methylation in the *Vinv* promoter region controls Cold Induced Sweetening in potato": Data files S1-S3

**Data File S1**

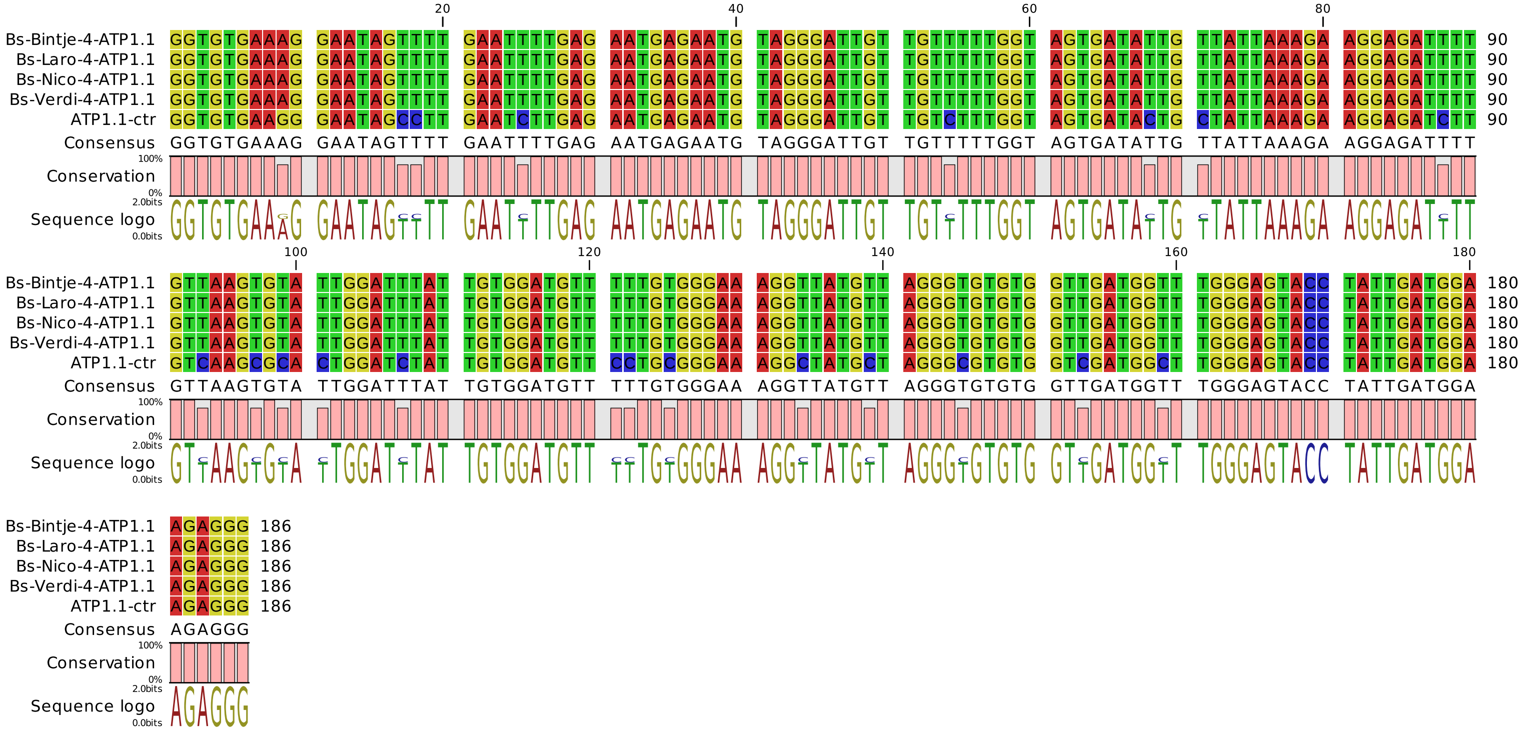


Alignment of ATP1.1 gene sequence from a non-bisulfite-treated representative sample (ATP1.1-ctr) and Bisulfite converted Bintje, Nico, Laro and Verdi after 4°C storage

**Data File S2**

### Alignment Bintje Vs Nico Vs Laro Vs Verdi

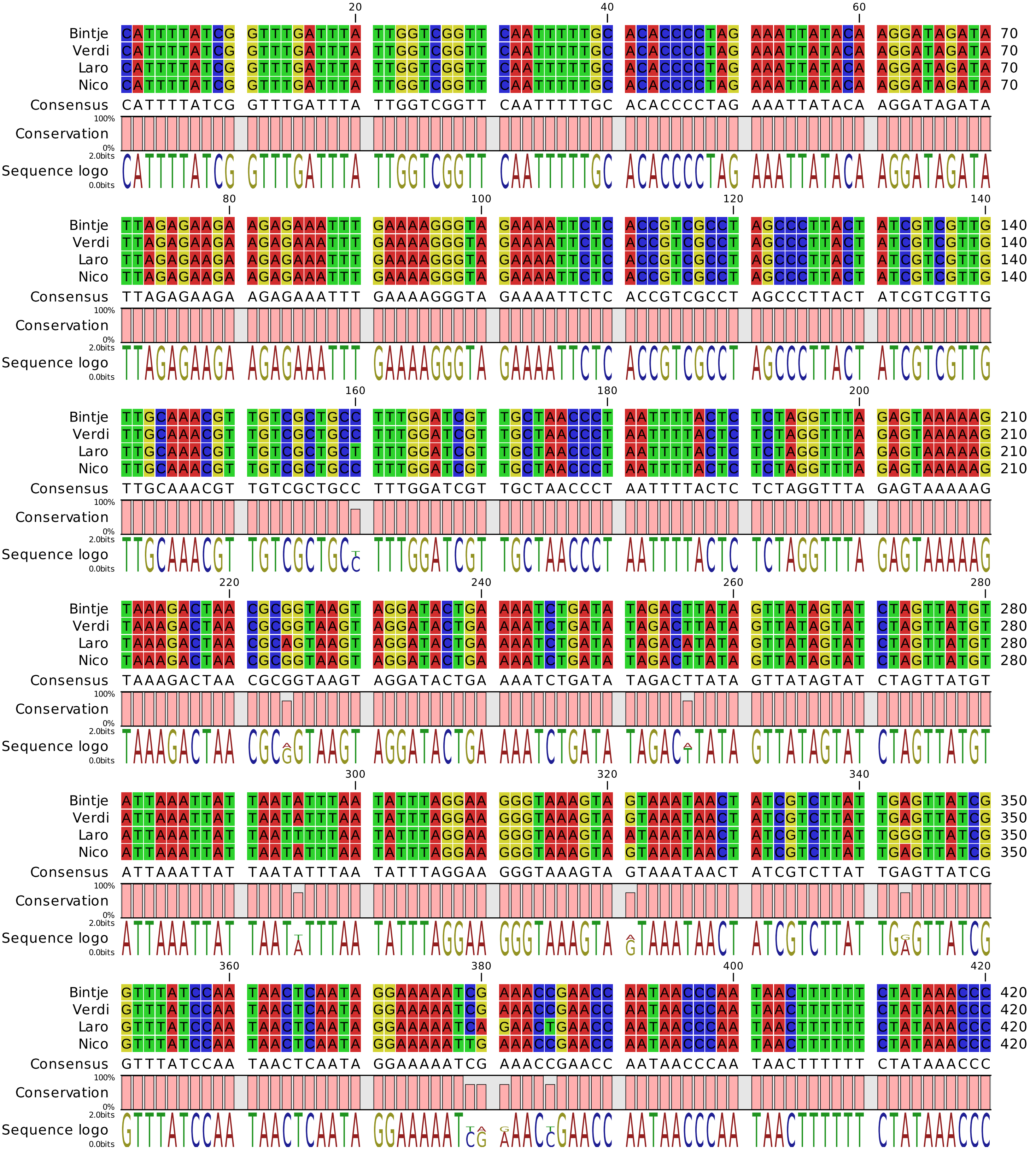

### Alignment Bintje Vs Nico Vs Laro Vs Verdi

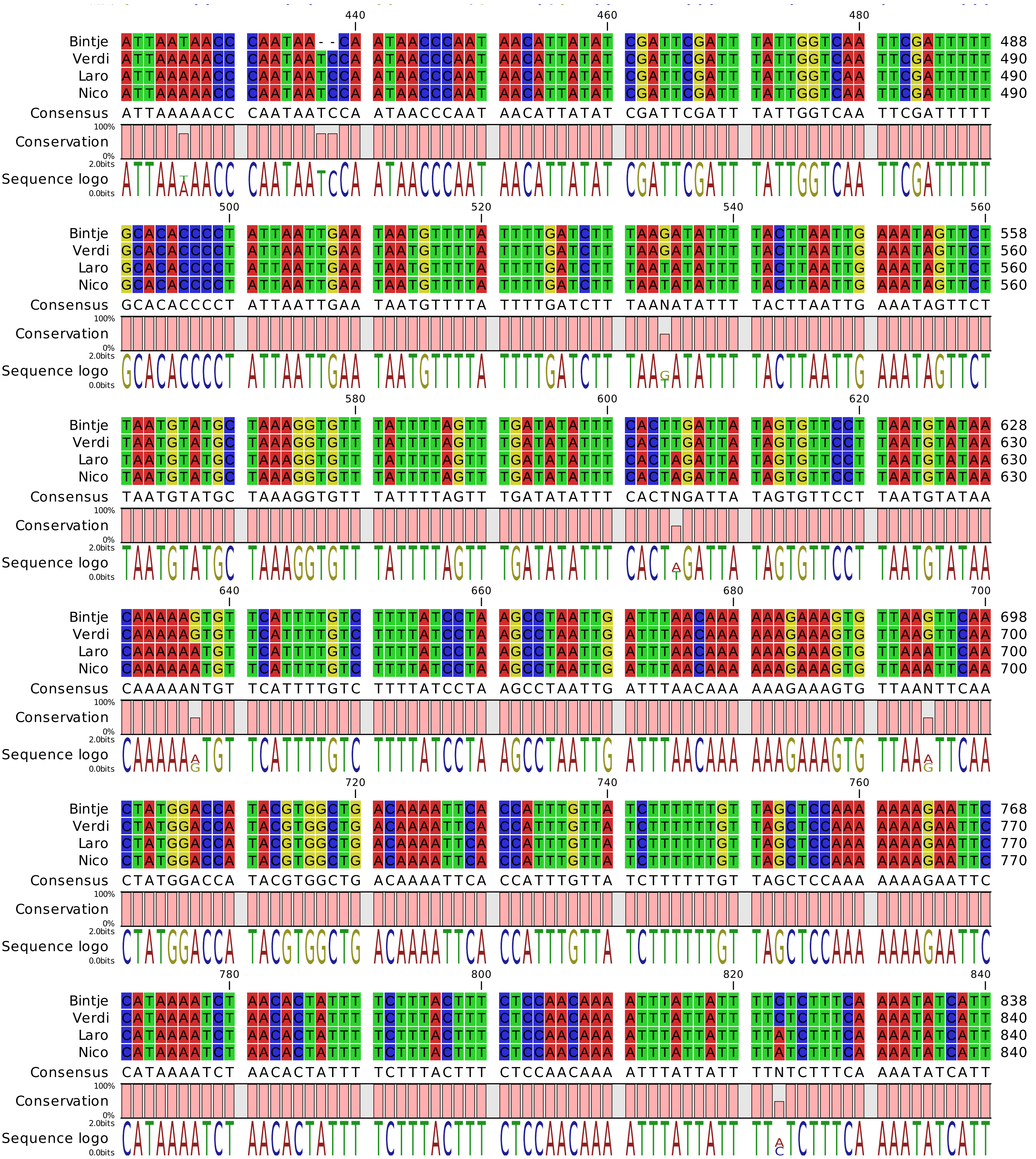

Alignment Bintje Vs Nico Vs Laro Vs Verdi

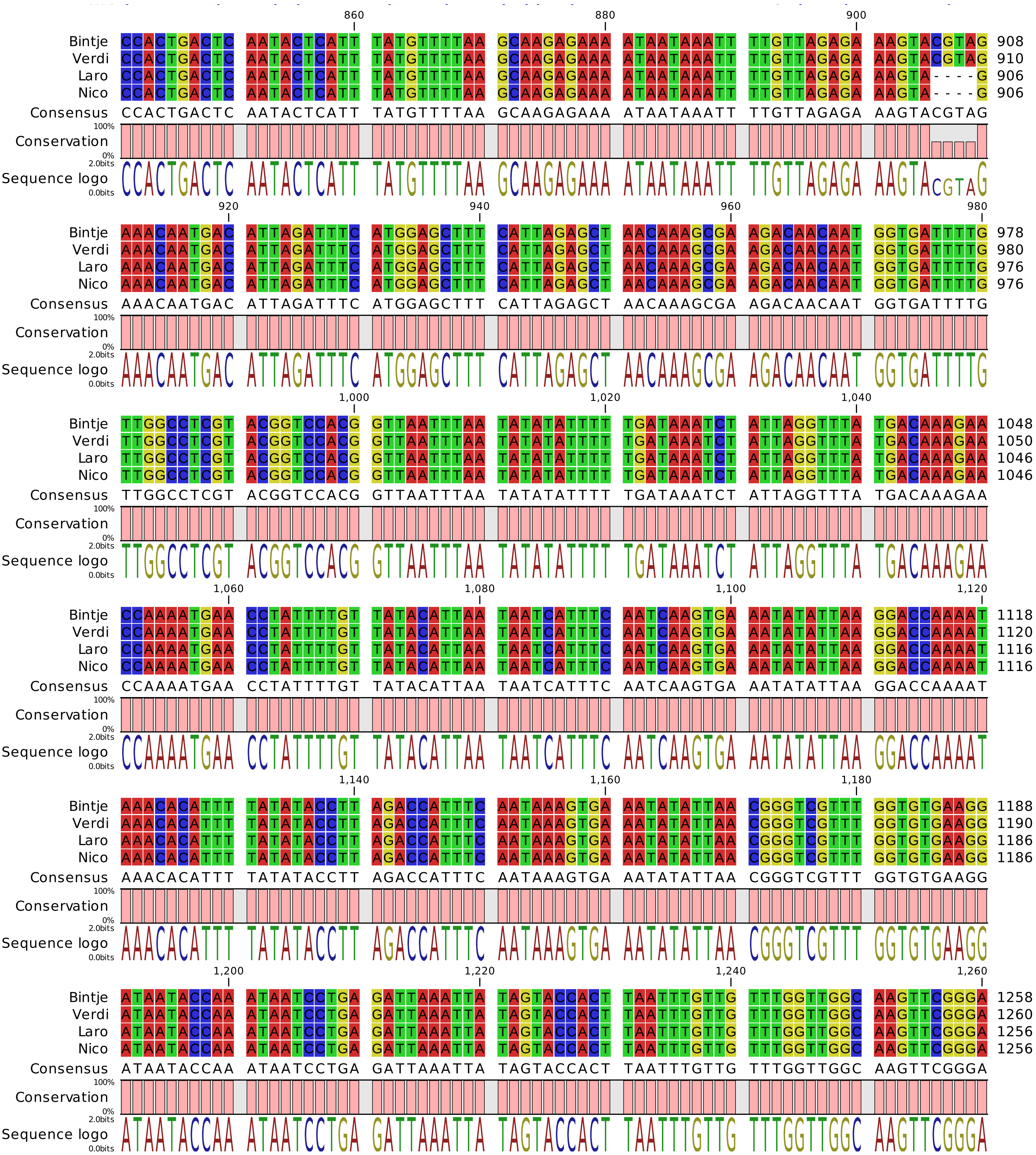

### Alignment Bintje Vs Nico Vs Laro Vs Verdi

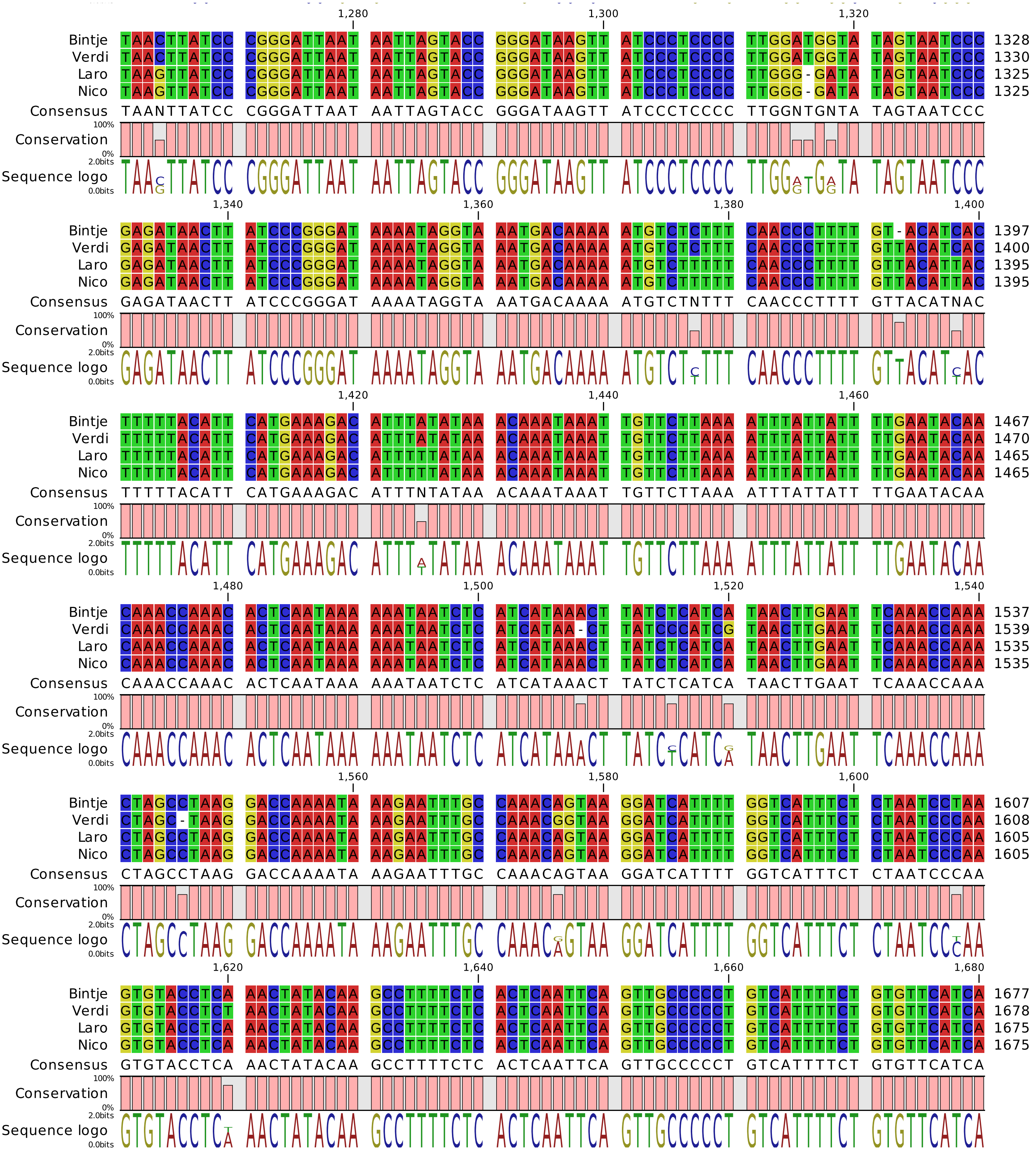

### Alignment Bintje Vs Nico Vs Laro Vs Verdi

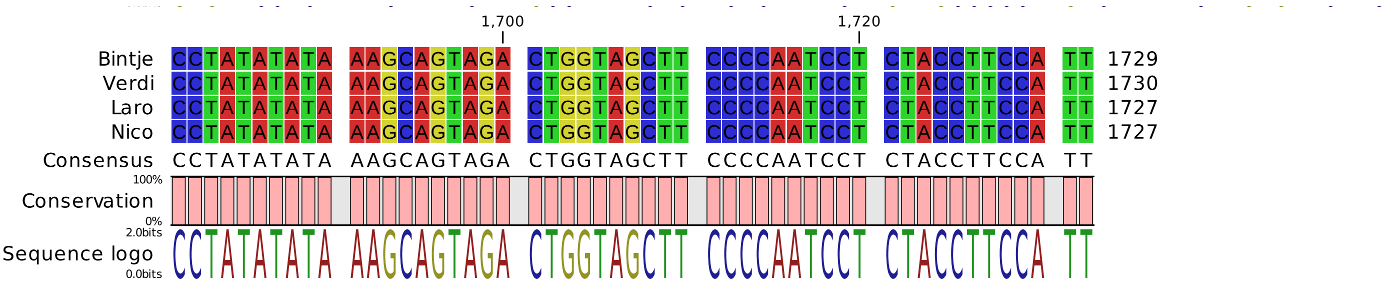

**Data File S3**

### Alignment Bintje Vs Bs-Bintje-4°

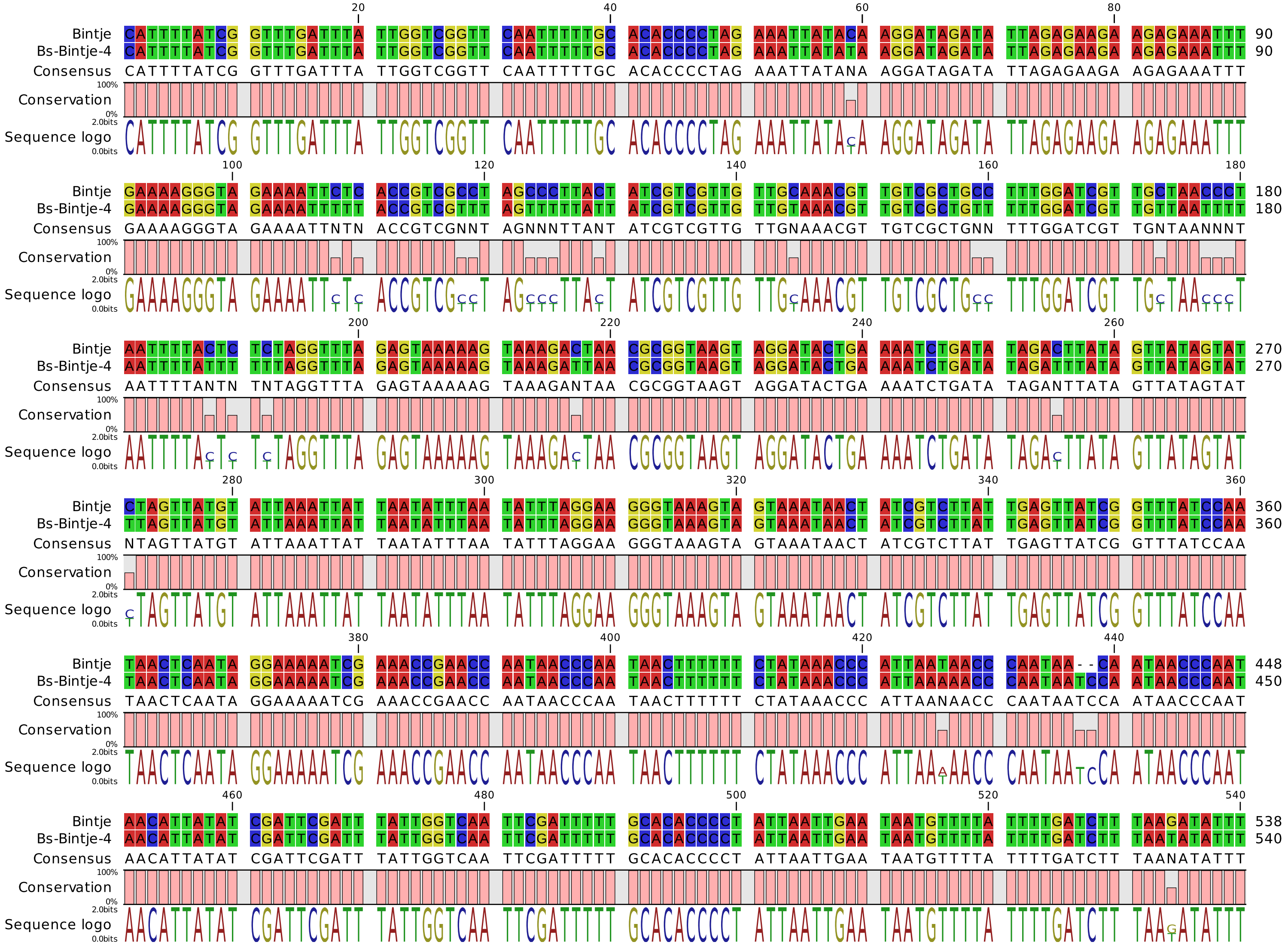

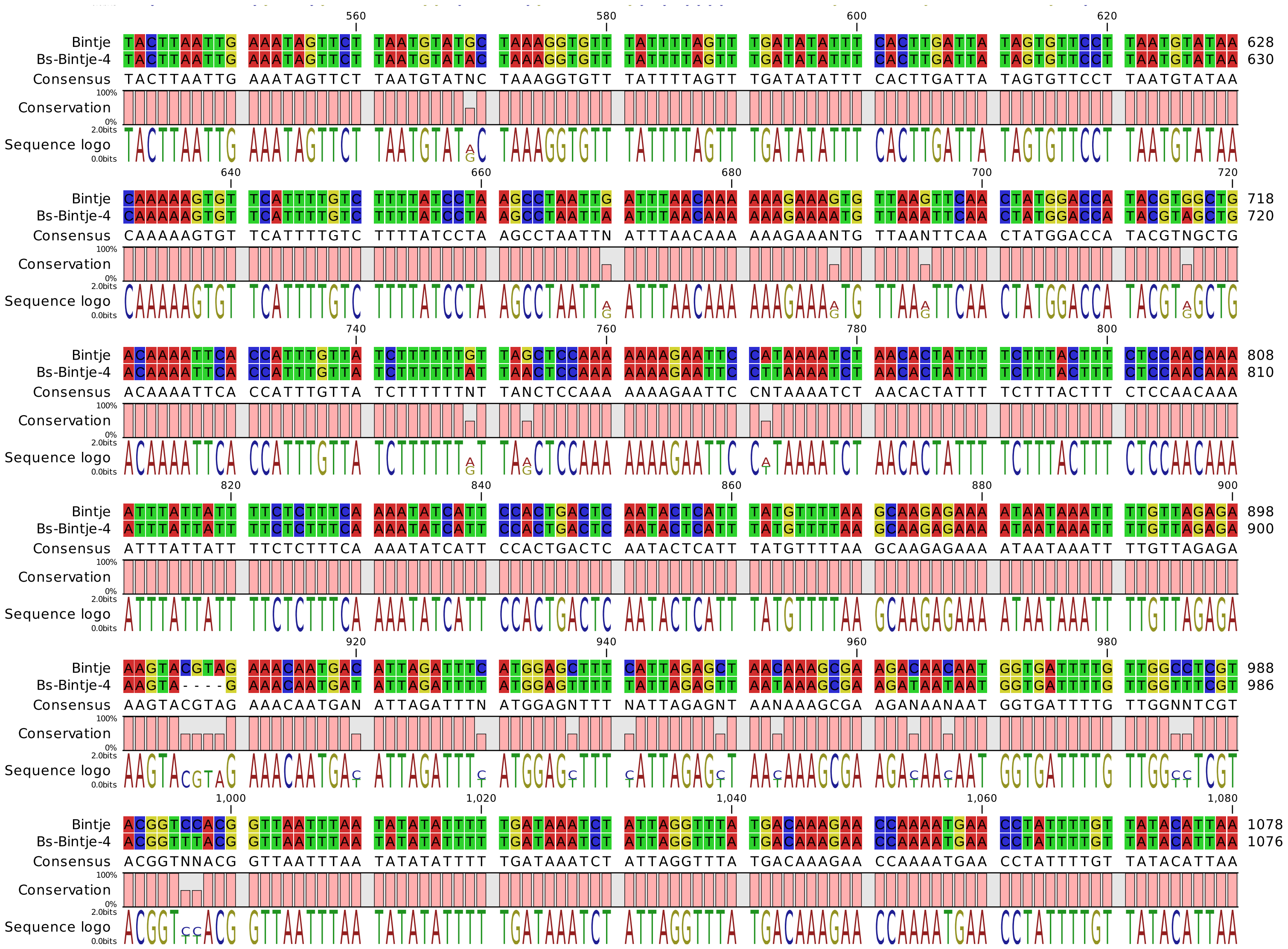

560 580 600 620
Bintje TACTTAATTG AAATAGTTCT TAATGTATGC TAAAGGTGTT TATTTTAGTT TGATATATTT CACTTGATTA TAGTGTTCCT TAATGTATAA 628
Bs-Bintje-4 TACTTAATTG AAATAGTTCT TAATGTATAC TAAAGGTGTT TATTTTAGTT TGATATATTT CACTTGATTA TAGTGTTCCT TAATGTATAA 630
Consensus TACTTAATTG AAATAGTTCT TAATGTATNC TAAAGGTGTT TATTTTAGTT TGATATATTT CACTTGATTA TAGTGTTCCT TAATGTATAA
Conservation
Sequence logo
640 660 680 700 720
Bintje CAAAAAGTGT TCATTTTGTC TTTTATCCTA AGCCTAATTG ATTTAACAAA AAAGAAAGTG TTAAGTTCAA CTATGGACCA TACGTGGCTG 718
Bs-Bintje-4 CAAAAAGTGT TCATTTTGTC TTTTATCCTA AGCCTAATTA ATTTAACAAA AAAGAAAATG TTAAATTCAA CTATGGACCA TACGTAGCTG 720
Consensus CAAAAAGTGT TCATTTTGTC TTTTATCCTA AGCCTAATTN ATTTAACAAA AAAGAAANTG TTAANTTCAA CTATGGACCA TACGTNGCTG
Conservation
Sequence logo
740 760 780 800
Bintje ACAAAATTCA CCATTTGTTA TCTTTTTTGT TAGCTCCAAA AAAAGAATTC CATAAAATCT AACACTATTT TCTTTACTTT CTCCAACAAA 808
Bs-Bintje-4 ACAAAATTCA CCATTTGTTA TCTTTTTTAT TAACTCCAAA AAAAGAATTC CTTAAAATCT AACACTATTT TCTTTACTTT CTCCAACAAA 810
Consensus ACAAAATTCA CCATTTGTTA TCTTTTTTNT TANCTCCAAA AAAAGAATTC CNTAAAATCT AACACTATTT TCTTTACTTT CTCCAACAAA
Conservation
Sequence logo
820 840 860 880 900
Bintje ATTTATTATT TTCTCTTTCA AAATATCATT CCACTGACTC AATACTCATT TATGTTTTAA GCAAGAGAAA ATAATAAATT TTGTTAGAGA 898
Bs-Bintje-4 ATTTATTATT TTCTCTTTCA AAATATCATT CCACTGACTC AATACTCATT TATGTTTTAA GCAAGAGAAA ATAATAAATT TTGTTAGAGA 900
Consensus ATTTATTATT TTCTCTTTCA AAATATCATT CCACTGACTC AATACTCATT TATGTTTTAA GCAAGAGAAA ATAATAAATT TTGTTAGAGA
Conservation
Sequence logo
920 940 960 980
Bintje AAGTACGTAG AAACAATGAC ATTAGATTTC ATGGAGCTTT CATTAGAGCT AACAAAGCGA AGACAACAAT GGTGATTTTG TTGGCCTCGT 988
Bs-Bintje-4 AAGTA----G AAACAATGAT ATTAGATTTT ATGGAGTTTT TATTAGAGTT AATAAAGCGA AGATAATAAT GGTGATTTTG TTGGTTTCGT 986
Consensus AAGTACGTAG AAACAATGAN ATTAGATTTN ATGGAGNTTT NATTAGAGNT AANAAAGCGA AGANAANAAT GGTGATTTTG TTGGNNTCGT
Conservation
Sequence logo
1,000 1,020 1,040 1,060 1,080
Bintje ACGGTCCACG GTTAATTTAA TATATATTTT TGATAAATCT ATTAGGTTTA TGACAAAGAA CCAAAATGAA CCTATTTTGT TATACATTAA 1078
Bs-Bintje-4 ACGGTTTACG GTTAATTTAA TATATATTTT TGATAAATCT ATTAGGTTTA TGACAAAGAA CCAAAATGAA CCTATTTTGT TATACATTAA 1076
Consensus ACGGTNNACG GTTAATTTAA TATATATTTT TGATAAATCT ATTAGGTTTA TGACAAAGAA CCAAAATGAA CCTATTTTGT TATACATTAA
Conservation
Sequence logo

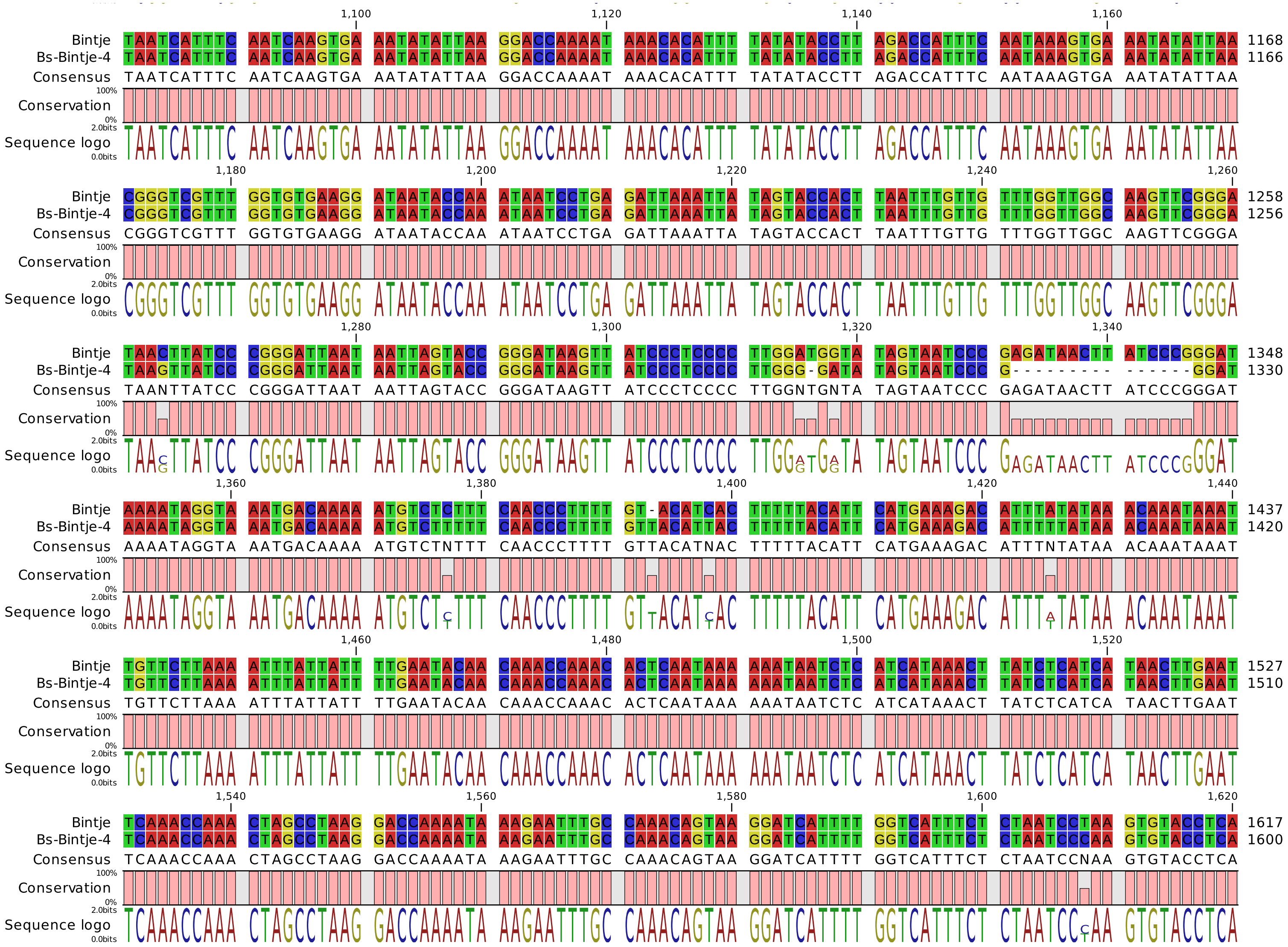
1,100 1,120 1,140 1,160
Bintje TAATCATTTC AATCAAGTGA AATATATTAA GGACCAAAAT AAACACATTT TATATACCTT AGACCATTTC AATAAAGTGA AATATATTAA 1168
Bs-Bintje-4 TAATCATTTC AATCAAGTGA AATATATTAA GGACCAAAAT AAACACATTT TATATACCTT AGACCATTTC AATAAAGTGA AATATATTAA 1166
Consensus TAATCATTTC AATCAAGTGA AATATATTAA GGACCAAAAT AAACACATTT TATATACCTT AGACCATTTC AATAAAGTGA AATATATTAA
Conservation
Sequence logo
1,180 1,200 1,220 1,240 1,260
Bintje CGGGTCGTTT GGTGTGAAGG ATAATACCAA ATAATCCTGA GATTAAATTA TAGTACCACT TAATTTGTTG TTTGGTTGGC AAGTTCGGGA 1258
Bs-Bintje-4 CGGGTCGTTT GGTGTGAAGG ATAATACCAA ATAATCCTGA GATTAAATTA TAGTACCACT TAATTTGTTG TTTGGTTGGC AAGTTCGGGA 1256
Consensus CGGGTCGTTT GGTGTGAAGG ATAATACCAA ATAATCCTGA GATTAAATTA TAGTACCACT TAATTTGTTG TTTGGTTGGC AAGTTCGGGA
Conservation
Sequence logo
1,280 1,300 1,320 1,340
Bintje TAACTTATCC CGGGATTAAT AATTAGTACC GGGATAAGTT ATCCCTCCCC TTGGATGGTA TAGTAATCCC GAGATAACTT ATCCCGGGAT 1348
Bs-Bintje-4 TAAGTTATCC CGGGATTAAT AATTAGTACC GGGATAAGTT ATCCCTCCCC TTGGG-GATA TAGTAATCCC G--------- ------GGAT 1330
Consensus TAANTTATCC CGGGATTAAT AATTAGTACC GGGATAAGTT ATCCCTCCCC TTGGNTGNTA TAGTAATCCC GAGATAACTT ATCCCGGGAT
Conservation
Sequence logo
1,360 1,380 1,400 1,420 1,440
Bintje AAAATAGGTA AATGACAAAA ATGTCTCTTT CAACCCTTTT GT-ACATCAC TTTTTACATT CATGAAAGAC ATTTATATAA ACAAATAAAT 1437
Bs-Bintje-4 AAAATAGGTA AATGACAAAA ATGTCTTTTT CAACCCTTTT GTTACATTAC TTTTTACATT CATGAAAGAC ATTTTTATAA ACAAATAAAT 1420
Consensus AAAATAGGTA AATGACAAAA ATGTCTNTTT CAACCCTTTT GTTACATNAC TTTTTACATT CATGAAAGAC ATTTNTATAA ACAAATAAAT
Conservation
Sequence logo
1,460 1,480 1,500 1,520
Bintje TGTTCTTAAA ATTTATTATT TTGAATACAA CAAACCAAAC ACTCAATAAA AAATAATCTC ATCATAAACT TATCTCATCA TAACTTGAAT 1527
Bs-Bintje-4 TGTTCTTAAA ATTTATTATT TTGAATACAA CAAACCAAAC ACTCAATAAA AAATAATCTC ATCATAAACT TATCTCATCA TAACTTGAAT 1510
Consensus TGTTCTTAAA ATTTATTATT TTGAATACAA CAAACCAAAC ACTCAATAAA AAATAATCTC ATCATAAACT TATCTCATCA TAACTTGAAT
Conservation
Sequence logo
1,540 1,560 1,580 1,600 1,620
Bintje TCAAACCAAA CTAGCCTAAG GACCAAAATA AAGAATTTGC CAAACAGTAA GGATCATTTT GGTCATTTCT CTAATCCTAA GTGTACCTCA 1617
Bs-Bintje-4 TCAAACCAAA CTAGCCTAAG GACCAAAATA AAGAATTTGC CAAACAGTAA GGATCATTTT GGTCATTTCT CTAATCCCAA GTGTACCTCA 1600
Consensus TCAAACCAAA CTAGCCTAAG GACCAAAATA AAGAATTTGC CAAACAGTAA GGATCATTTT GGTCATTTCT CTAATCCNAA GTGTACCTCA
Conservation
Sequence logo

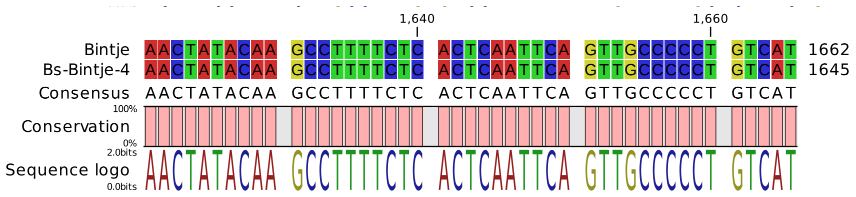
1,640
1,660
Bintje AACTATACAA GCCTTTTCTC ACTCAATTCA GTTGCCCCCT GTCAT 1662
Bs-Bintje-4 AACTATACAA GCCTTTTCTC ACTCAATTCA GTTGCCCCCT GTCAT 1645
Consensus AACTATACAA GCCTTTTCTC ACTCAATTCA GTTGCCCCCT GTCAT
100%
Conservation
0%
2.0bits
Sequence logo
0.0bits

### Alignment Laro Vs Bs-Laro-4°

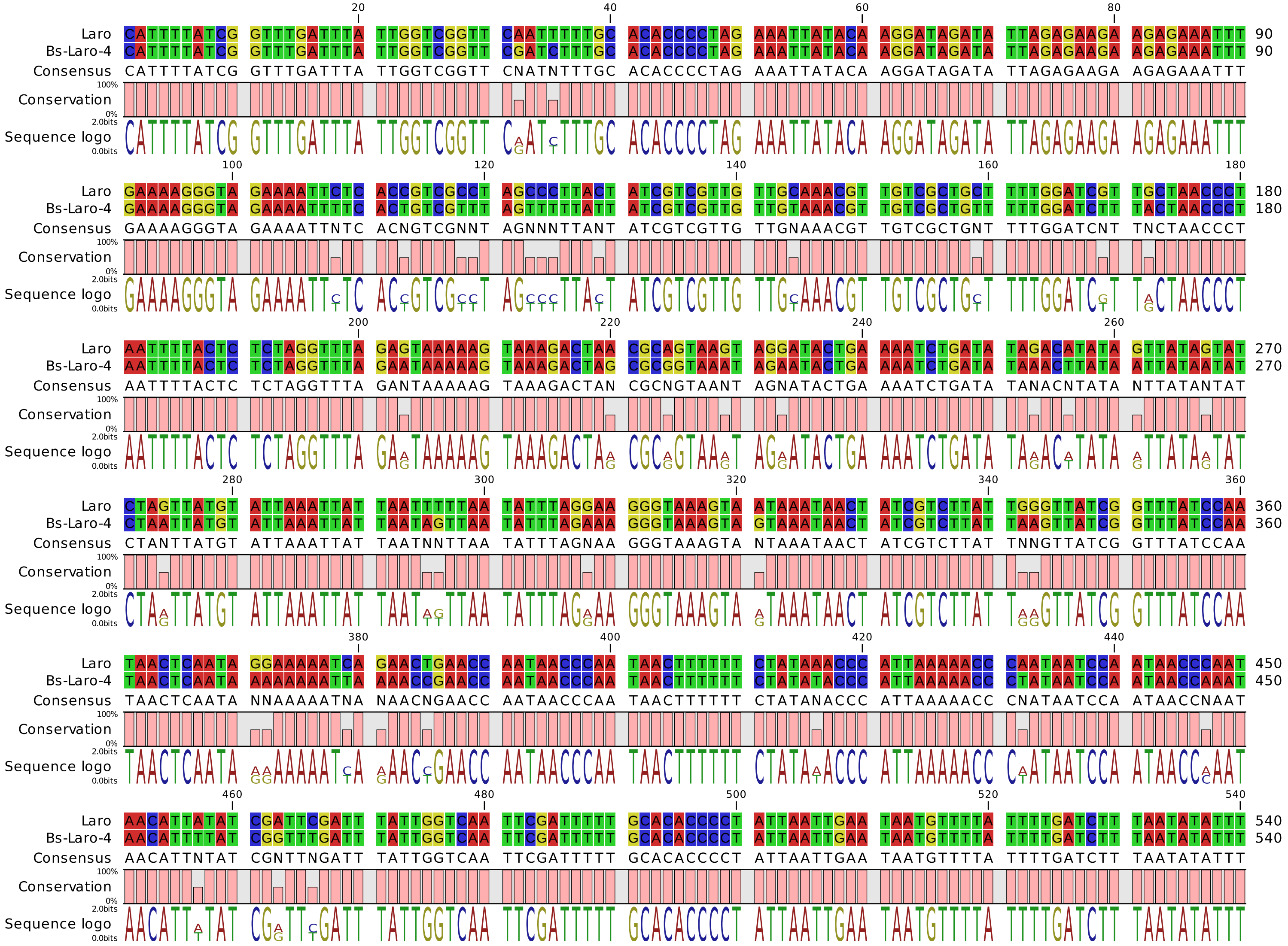

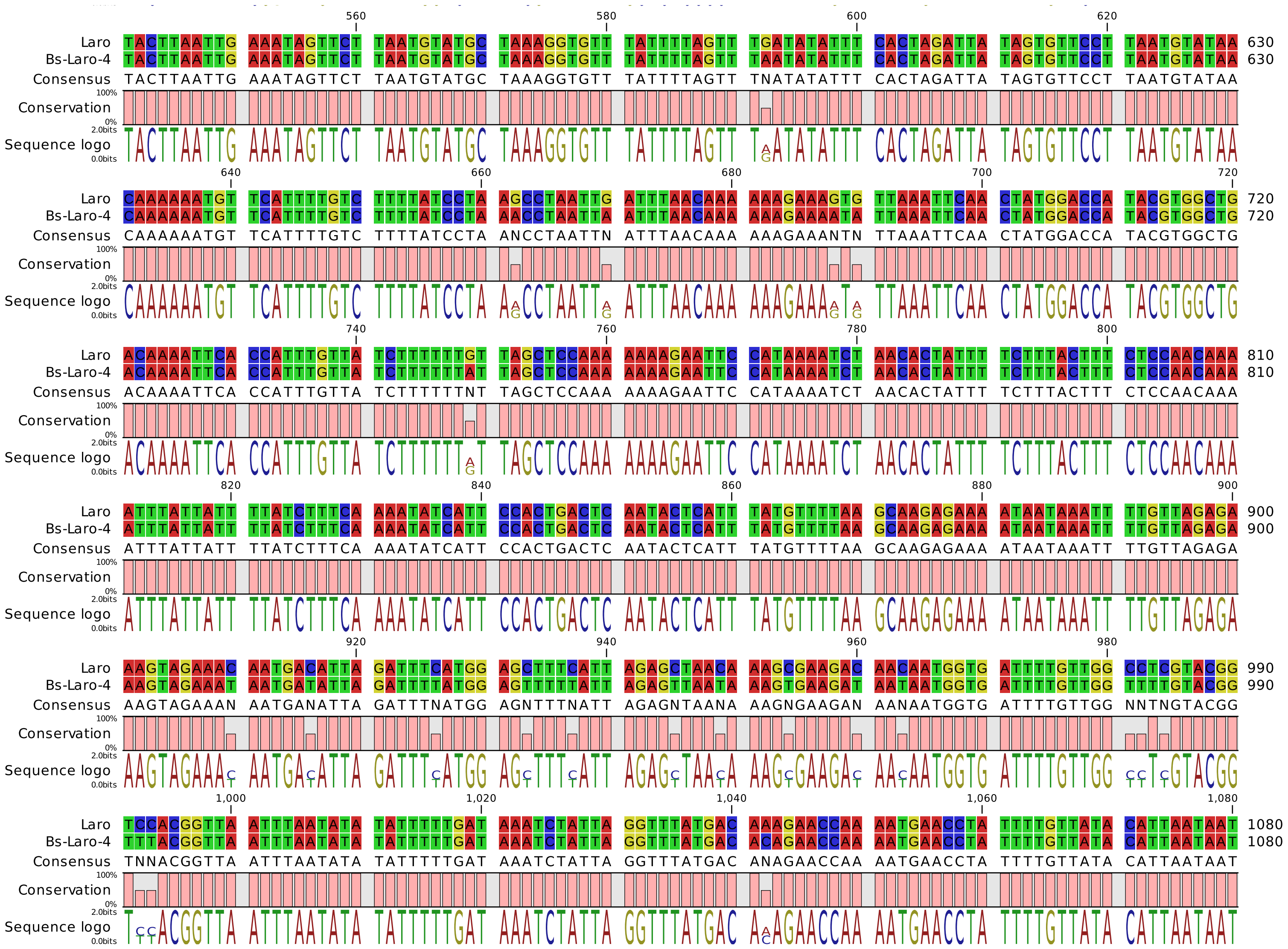
Laro TACTTAATTG AAATAGTTCT TAATGTATGC TAAAGGTGTT TATTTTAGTT TGATATATTT CACTAGATTA TAGTGTTCCT TAATGTATAA 630
Bs-Laro-4 TACTTAATTG AAATAGTTCT TAATGTATGC TAAAGGTGTT TATTTTAGTT TAATATATTT CACTAGATTA TAGTGTTCCT TAATGTATAA 630
Consensus TACTTAATTG AAATAGTTCT TAATGTATGC TAAAGGTGTT TATTTTAGTT TNATATATTT CACTAGATTA TAGTGTTCCT TAATGTATAA
Conservation
Sequence logo
Laro CAAAAAATGT TCATTTTGTC TTTTATCCTA AGCCTAATTG ATTTAACAAA AAAGAAAGTG TTAAATTCAA CTATGGACCA TACGTGGCTG 720
Bs-Laro-4 CAAAAAATGT TCATTTTGTC TTTTATCCTA AACCTAATTA ATTTAACAAA AAAGAAAATA TTAAATTCAA CTATGGACCA TACGTGGCTG 720
Consensus CAAAAAATGT TCATTTTGTC TTTTATCCTA ANCCTAATTN ATTTAACAAA AAAGAAANTN TTAAATTCAA CTATGGACCA TACGTGGCTG
Laro ACAAAATTCA CCATTTGTTA TCTTTTTTGT TAGCTCCAAA AAAAGAATTC CATAAAATCT AACACTATTT TCTTTACTTT CTCCAACAAA 810
Bs-Laro-4 ACAAAATTCA CCATTTGTTA TCTTTTTTAT TAGCTCCAAA AAAAGAATTC CATAAAATCT AACACTATTT TCTTTACTTT CTCCAACAAA 810
Consensus ACAAAATTCA CCATTTGTTA TCTTTTTTNT TAGCTCCAAA AAAAGAATTC CATAAAATCT AACACTATTT TCTTTACTTT CTCCAACAAA
Laro ATTTATTATT TTATCTTTCA AAATATCATT CCACTGACTC AATACTCATT TATGTTTTAA GCAAGAGAAA ATAATAAATT TTGTTAGAGA 900
Bs-Laro-4 ATTTATTATT TTATCTTTCA AAATATCATT CCACTGACTC AATACTCATT TATGTTTTAA GCAAGAGAAA ATAATAAATT TTGTTAGAGA 900
Consensus ATTTATTATT TTATCTTTCA AAATATCATT CCACTGACTC AATACTCATT TATGTTTTAA GCAAGAGAAA ATAATAAATT TTGTTAGAGA
Laro AAGTAGAAAC AATGACATTA GATTTCATGG AGCTTTCATT AGAGCTAACA AAGCGAAGAC AACAATGGTG ATTTTGTTGG CCTCGTACGG 990
Bs-Laro-4 AAGTAGAAAT AATGATATTA GATTTTATGG AGTTTTTATT AGAGTTAATA AAGTGAAGAT AATAATGGTG ATTTTGTTGG TTTTGTACGG 990
Consensus AAGTAGAAAN AATGANATTA GATTTNATGG AGNTTTNATT AGAGNTAANA AAGNGAAGAN AANAATGGTG ATTTTGTTGG NNTNGTACGG
Laro TCCACGGTTA ATTTAATATA TATTTTTGAT AAATCTATTA GGTTTATGAC AAAGAACCAA AATGAACCTA TTTTGTTATA CATTAATAAT 1080
Bs-Laro-4 TTTACGGTTA ATTTAATATA TATTTTTGAT AAATCTATTA GGTTTATGAC ACAGAACCAA AATGAACCTA TTTTGTTATA CATTAATAAT 1080
Consensus TNNACGGTTA ATTTAATATA TATTTTTGAT AAATCTATTA GGTTTATGAC ANAGAACCAA AATGAACCTA TTTTGTTATA CATTAATAAT

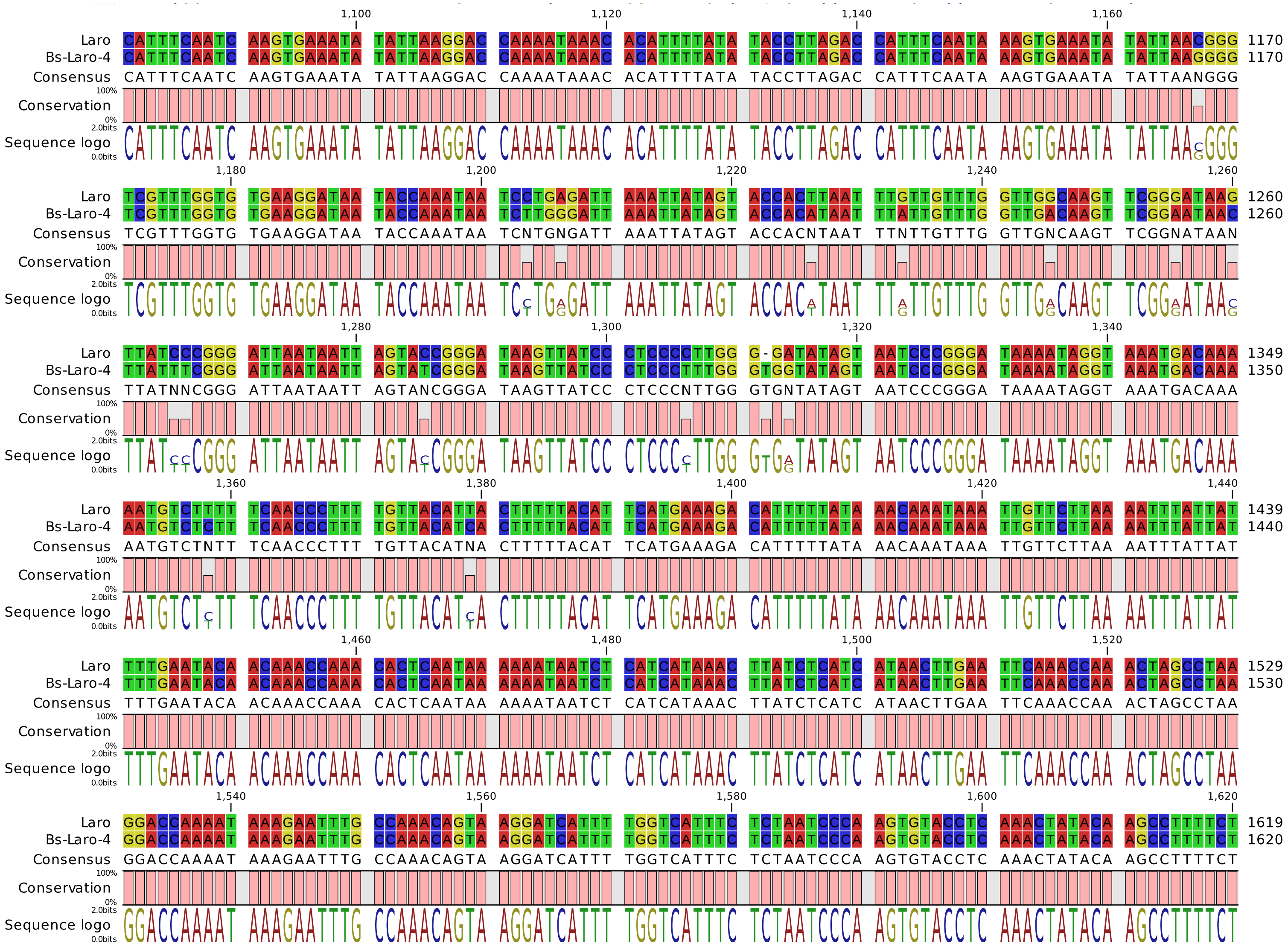
1,100 1,120 1,140 1,160
Laro CATTTCAATC AAGTGAAATA TATTAAGGAC CAAAATAAAC ACATTTTATA TACCTTAGAC CATTTCAATA AAGTGAAATA TATTAACGGG 1170
Bs-Laro-4 CATTTCAATC AAGTGAAATA TATTAAGGAC CAAAATAAAC ACATTTTATA TACCTTAGAC CATTTCAATA AAGTGAAATA TATTAAGGGG 1170
Consensus CATTTCAATC AAGTGAAATA TATTAAGGAC CAAAATAAAC ACATTTTATA TACCTTAGAC CATTTCAATA AAGTGAAATA TATTAANGGG
Conservation
Sequence logo
1,180 1,200 1,220 1,240 1,260
Laro TCGTTTGGTG TGAAGGATAA TACCAAATAA TCCTGAGATT AAATTATAGT ACCACTTAAT TTGTTGTTTG GTTGGCAAGT TCGGGATAAG 1260
Bs-Laro-4 TCGTTTGGTG TGAAGGATAA TACCAAATAA TCTTGGGATT AAATTATAGT ACCACATAAT TTATTGTTTG GTTGACAAGT TCGGAATAAC 1260
Consensus TCGTTTGGTG TGAAGGATAA TACCAAATAA TCNTGNGATT AAATTATAGT ACCACNTAAT TTNTTGTTTG GTTGNCAAGT TCGGNATAAN
Conservation
Sequence logo
1,280 1,300 1,320 1,340
Laro TTATCCCGGG ATTAATAATT AGTACCGGGA TAAGTTATCC CTCCCCTTGG G-GATATAGT AATCCCGGGA TAAAATAGGT AAATGACAAA 1349
Bs-Laro-4 TTATTTCGGG ATTAATAATT AGTATCGGGA TAAGTTATCC CTCCCTTTGG GTGGTATAGT AATCCCGGGA TAAAATAGGT AAATGACAAA 1350
Consensus TTATNNCGGG ATTAATAATT AGTANCGGGA TAAGTTATCC CTCCCNTTGG GTGNTATAGT AATCCCGGGA TAAAATAGGT AAATGACAAA
Conservation
Sequence logo
1,360 1,380 1,400 1,420 1,440
Laro AATGTCTTTT TCAACCCTTT TGTTACATTA CTTTTTACAT TCATGAAAGA CATTTTTATA AACAAATAAA TTGTTCTTAA AATTTATTAT 1439
Bs-Laro-4 AATGTCTCTT TCAACCCTTT TGTTACATCA CTTTTTACAT TCATGAAAGA CATTTTTATA AACAAATAAA TTGTTCTTAA AATTTATTAT 1440
Consensus AATGTCTNTT TCAACCCTTT TGTTACATNA CTTTTTACAT TCATGAAAGA CATTTTTATA AACAAATAAA TTGTTCTTAA AATTTATTAT
Conservation
Sequence logo
1,460 1,480 1,500 1,520
Laro TTTGAATACA ACAAACCAAA CACTCAATAA AAAATAATCT CATCATAAAC TTATCTCATC ATAACTTGAA TTCAAACCAA ACTAGCCTAA 1529
Bs-Laro-4 TTTGAATACA ACAAACCAAA CACTCAATAA AAAATAATCT CATCATAAAC TTATCTCATC ATAACTTGAA TTCAAACCAA ACTAGCCTAA 1530
Consensus TTTGAATACA ACAAACCAAA CACTCAATAA AAAATAATCT CATCATAAAC TTATCTCATC ATAACTTGAA TTCAAACCAA ACTAGCCTAA
Conservation
Sequence logo
1,540 1,560 1,580 1,600 1,620
Laro GGACCAAAAT AAAGAATTTG CCAAACAGTA AGGATCATTT TGGTCATTTC TCTAATCCCA AGTGTACCTC AAACTATACA AGCCTTTTCT 1619
Bs-Laro-4 GGACCAAAAT AAAGAATTTG CCAAACAGTA AGGATCATTT TGGTCATTTC TCTAATCCCA AGTGTACCTC AAACTATACA AGCCTTTTCT 1620
Consensus GGACCAAAAT AAAGAATTTG CCAAACAGTA AGGATCATTT TGGTCATTTC TCTAATCCCA AGTGTACCTC AAACTATACA AGCCTTTTCT
Conservation
Sequence logo

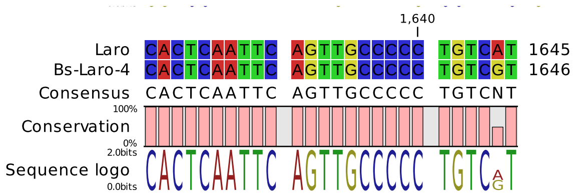
1,640
Laro CACTCAATTC AGTTGCCCCC TGTCAT 1645
Bs-Laro-4 CACTCAATTC AGTTGCCCCC TGTCGT 1646
Consensus CACTCAATTC AGTTGCCCCC TGTCNT
100%
Conservation
0%
2.0bits
Sequence logo
0.0bits

### Alignment Nico Vs Bs-Nico-4°

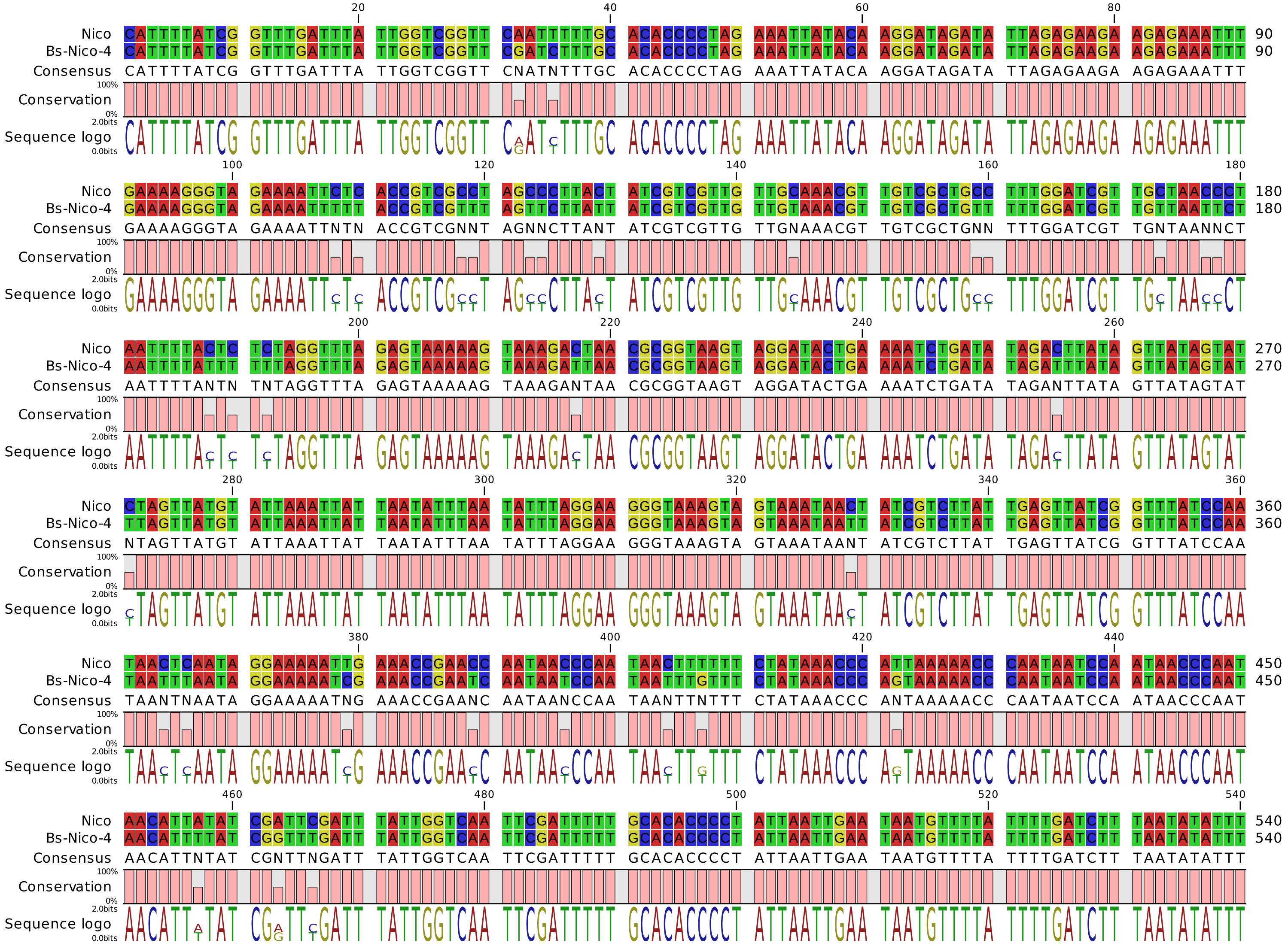

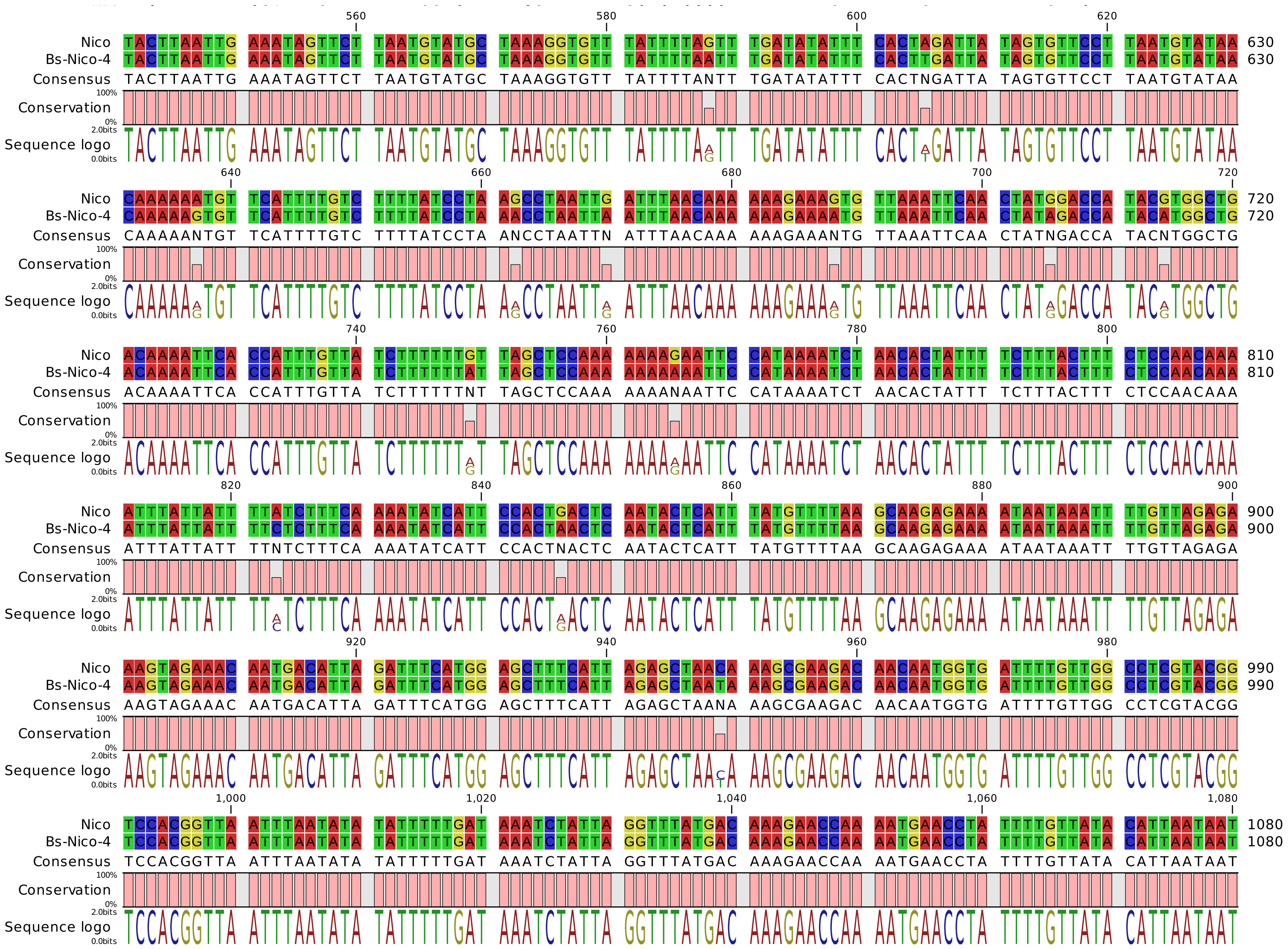
560 580 600 620
Nico TACTTAATTG AAATAGTTCT TAATGTATGC TAAAGGTGTT TATTTTAGTT TGATATATTT CACTAGATTA TAGTGTTCCT TAATGTATAA 630
Bs-Nico-4 TACTTAATTG AAATAGTTCT TAATGTATGC TAAAGGTGTT TATTTTAATT TGATATATTT CACTTGATTA TAGTGTTCCT TAATGTATAA 630
Consensus TACTTAATTG AAATAGTTCT TAATGTATGC TAAAGGTGTT TATTTTANTT TGATATATTT CACTNGATTA TAGTGTTCCT TAATGTATAA
Conservation
Sequence logo
640 660 680 700 720
Nico CAAAAAATGT TCATTTTGTC TTTTATCCTA AGCCTAATTG ATTTAACAAA AAAGAAAGTG TTAAATTCAA CTATGGACCA TACGTGGCTG 720
Bs-Nico-4 CAAAAAGTGT TCATTTTGTC TTTTATCCTA AACCTAATTA ATTTAACAAA AAAGAAAATG TTAAATTCAA CTATAGACCA TACATGGCTG 720
Consensus CAAAAANTGT TCATTTTGTC TTTTATCCTA ANCCTAATTN ATTTAACAAA AAAGAAANTG TTAAATTCAA CTATNGACCA TACNTGGCTG
Conservation
Sequence logo
740 760 780 800
Nico ACAAAATTCA CCATTTGTTA TCTTTTTTGT TAGCTCCAAA AAAAGAATTC CATAAAATCT AACACTATTT TCTTTACTTT CTCCAACAAA 810
Bs-Nico-4 ACAAAATTCA CCATTTGTTA TCTTTTTTAT TAGCTCCAAA AAAAAAATTC CATAAAATCT AACACTATTT TCTTTACTTT CTCCAACAAA 810
Consensus ACAAAATTCA CCATTTGTTA TCTTTTTTNT TAGCTCCAAA AAAANAATTC CATAAAATCT AACACTATTT TCTTTACTTT CTCCAACAAA
Conservation
Sequence logo
820 840 860 880 900
Nico ATTTATTATT TTATCTTTCA AAATATCATT CCACTGACTC AATACTCATT TATGTTTTAA GCAAGAGAAA ATAATAAATT TTGTTAGAGA 900
Bs-Nico-4 ATTTATTATT TTCTCTTTCA AAATATCATT CCACTAACTC AATACTCATT TATGTTTTAA GCAAGAGAAA ATAATAAATT TTGTTAGAGA 900
Consensus ATTTATTATT TTNTCTTTCA AAATATCATT CCACTNACTC AATACTCATT TATGTTTTAA GCAAGAGAAA ATAATAAATT TTGTTAGAGA
Conservation
Sequence logo
920 940 960 980
Nico AAGTAGAAAC AATGACATTA GATTTCATGG AGCTTTCATT AGAGCTAACA AAGCGAAGAC AACAATGGTG ATTTTGTTGG CCTCGTACGG 990
Bs-Nico-4 AAGTAGAAAC AATGACATTA GATTTCATGG AGCTTTCATT AGAGCTAATA AAGCGAAGAC AACAATGGTG ATTTTGTTGG CCTCGTACGG 990
Consensus AAGTAGAAAC AATGACATTA GATTTCATGG AGCTTTCATT AGAGCTAANA AAGCGAAGAC AACAATGGTG ATTTTGTTGG CCTCGTACGG
Conservation
Sequence logo
1,000 1,020 1,040 1,060 1,080
Nico TCCACGGTTA ATTTAATATA TATTTTTGAT AAATCTATTA GGTTTATGAC AAAGAACCAA AATGAACCTA TTTTGTTATA CATTAATAAT 1080
Bs-Nico-4 TCCACGGTTA ATTTAATATA TATTTTTGAT AAATCTATTA GGTTTATGAC AAAGAACCAA AATGAACCTA TTTTGTTATA CATTAATAAT 1080
Consensus TCCACGGTTA ATTTAATATA TATTTTTGAT AAATCTATTA GGTTTATGAC AAAGAACCAA AATGAACCTA TTTTGTTATA CATTAATAAT
Conservation
Sequence logo

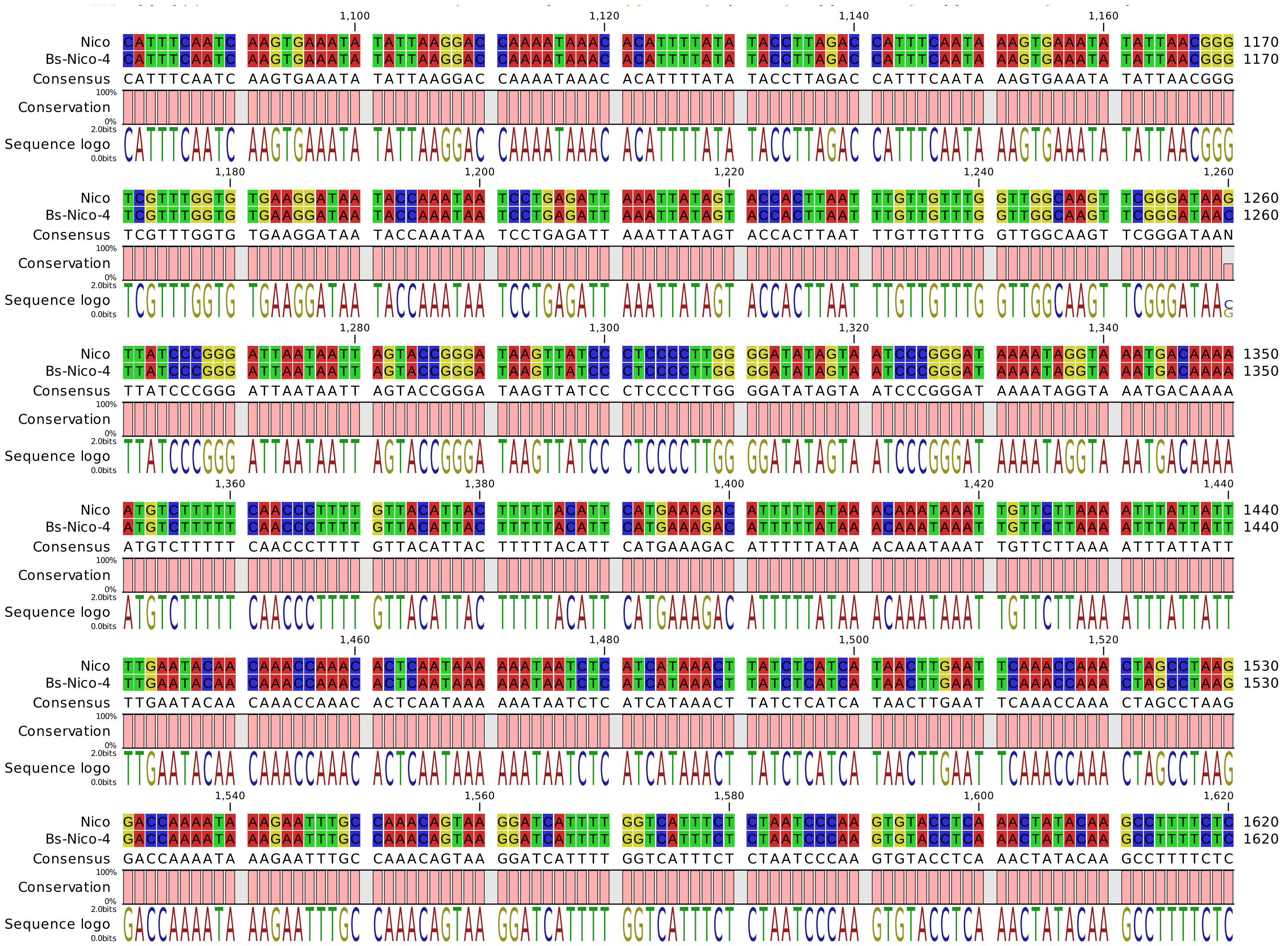

1,100 1,120 1,140 1,160
Nico CATTTCAATC AAGTGAAATA TATTAAGGAC CAAAATAAAC ACATTTTATA TACCTTAGAC CATTTCAATA AAGTGAAATA TATTAACGGG 1170
Bs-Nico-4 CATTTCAATC AAGTGAAATA TATTAAGGAC CAAAATAAAC ACATTTTATA TACCTTAGAC CATTTCAATA AAGTGAAATA TATTAACGGG 1170
Consensus CATTTCAATC AAGTGAAATA TATTAAGGAC CAAAATAAAC ACATTTTATA TACCTTAGAC CATTTCAATA AAGTGAAATA TATTAACGGG
Conservation
Sequence logo
1,180 1,200 1,220 1,240 1,260
Nico TCGTTTGGTG TGAAGGATAA TACCAAATAA TCCTGAGATT AAATTATAGT ACCACTTAAT TTGTTGTTTG GTTGGCAAGT TCGGGATAAG 1260
Bs-Nico-4 TCGTTTGGTG TGAAGGATAA TACCAAATAA TCCTGAGATT AAATTATAGT ACCACTTAAT TTGTTGTTTG GTTGGCAAGT TCGGGATAAC 1260
Consensus TCGTTTGGTG TGAAGGATAA TACCAAATAA TCCTGAGATT AAATTATAGT ACCACTTAAT TTGTTGTTTG GTTGGCAAGT TCGGGATAAN
Conservation
Sequence logo
1,280 1,300 1,320 1,340
Nico TTATCCCGGG ATTAATAATT AGTACCGGGA TAAGTTATCC CTCCCCTTGG GGATATAGTA ATCCCGGGAT AAAATAGGTA AATGACAAAA 1350
Bs-Nico-4 TTATCCCGGG ATTAATAATT AGTACCGGGA TAAGTTATCC CTCCCCTTGG GGATATAGTA ATCCCGGGAT AAAATAGGTA AATGACAAAA 1350
Consensus TTATCCCGGG ATTAATAATT AGTACCGGGA TAAGTTATCC CTCCCCTTGG GGATATAGTA ATCCCGGGAT AAAATAGGTA AATGACAAAA
Conservation
Sequence logo
1,360 1,380 1,400 1,420 1,440
Nico ATGTCTTTTT CAACCCTTTT GTTACATTAC TTTTTACATT CATGAAAGAC ATTTTTATAA ACAAATAAAT TGTTCTTAAA ATTTATTATT 1440
Bs-Nico-4 ATGTCTTTTT CAACCCTTTT GTTACATTAC TTTTTACATT CATGAAAGAC ATTTTTATAA ACAAATAAAT TGTTCTTAAA ATTTATTATT 1440
Consensus ATGTCTTTTT CAACCCTTTT GTTACATTAC TTTTTACATT CATGAAAGAC ATTTTTATAA ACAAATAAAT TGTTCTTAAA ATTTATTATT
Conservation
Sequence logo
1,460 1,480 1,500 1,520
Nico TTGAATACAA CAAACCAAAC ACTCAATAAA AAATAATCTC ATCATAAACT TATCTCATCA TAACTTGAAT TCAAACCAAA CTAGCCTAAG 1530
Bs-Nico-4 TTGAATACAA CAAACCAAAC ACTCAATAAA AAATAATCTC ATCATAAACT TATCTCATCA TAACTTGAAT TCAAACCAAA CTAGCCTAAG 1530
Consensus TTGAATACAA CAAACCAAAC ACTCAATAAA AAATAATCTC ATCATAAACT TATCTCATCA TAACTTGAAT TCAAACCAAA CTAGCCTAAG
Conservation
Sequence logo
1,540 1,560 1,580 1,600 1,620
Nico GACCAAAATA AAGAATTTGC CAAACAGTAA GGATCATTTT GGTCATTTCT CTAATCCCAA GTGTACCTCA AACTATACAA GCCTTTTCTC 1620
Bs-Nico-4 GACCAAAATA AAGAATTTGC CAAACAGTAA GGATCATTTT GGTCATTTCT CTAATCCCAA GTGTACCTCA AACTATACAA GCCTTTTCTC 1620
Consensus GACCAAAATA AAGAATTTGC CAAACAGTAA GGATCATTTT GGTCATTTCT CTAATCCCAA GTGTACCTCA AACTATACAA GCCTTTTCTC
Conservation
Sequence logo

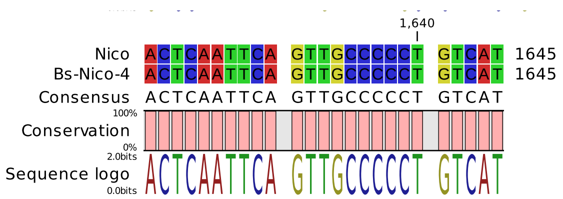
1,640
Nico ACTCAATTCA GTTGCCCCCT GTCAT 1645
Bs-Nico-4 ACTCAATTCA GTTGCCCCCT GTCAT 1645
Consensus ACTCAATTCA GTTGCCCCCT GTCAT
Conservation
100%
0%
Sequence logo
2.0bits
0.0bits

### Alignment Verdi Vs Bs-Verdi-4°

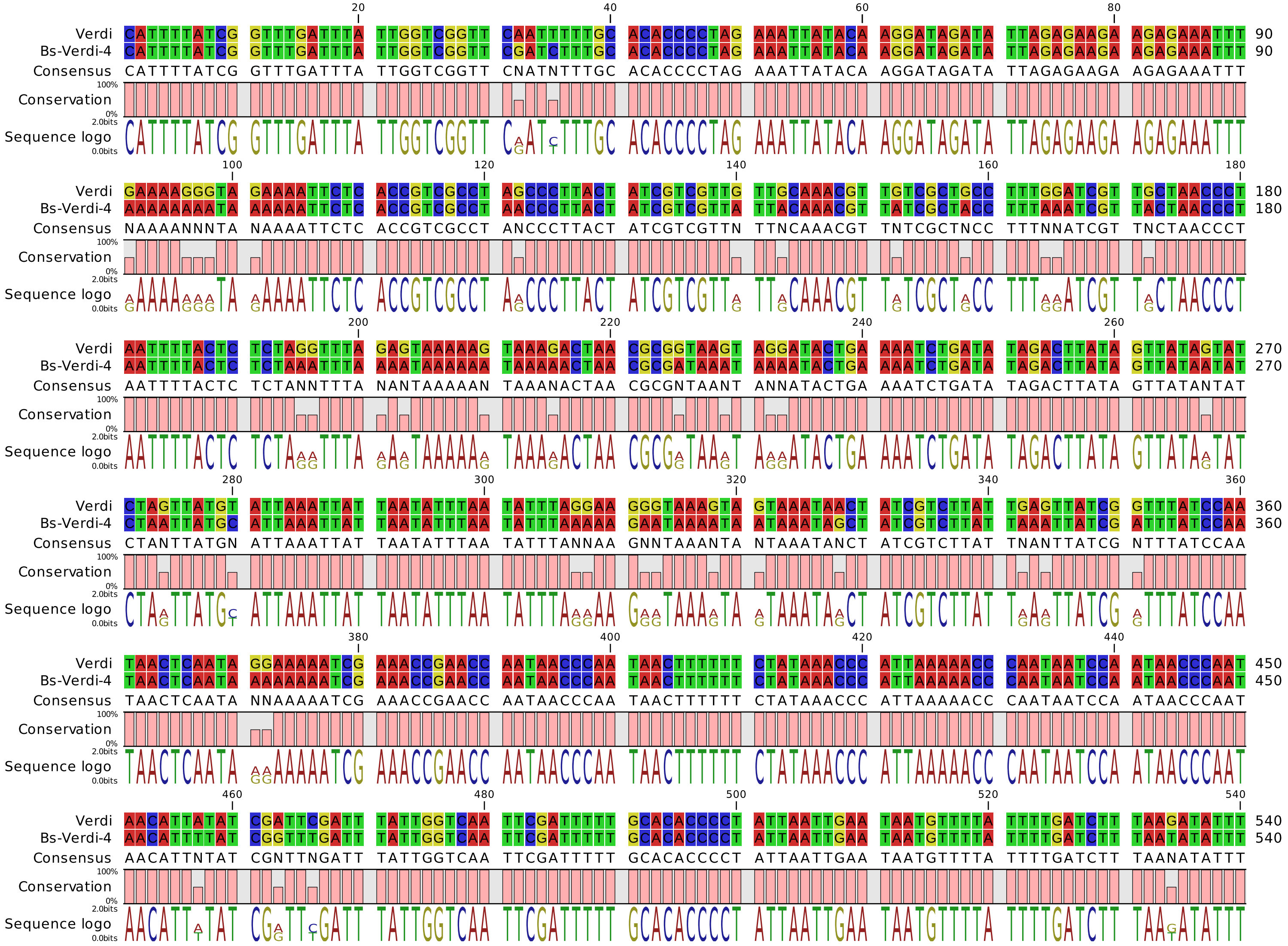

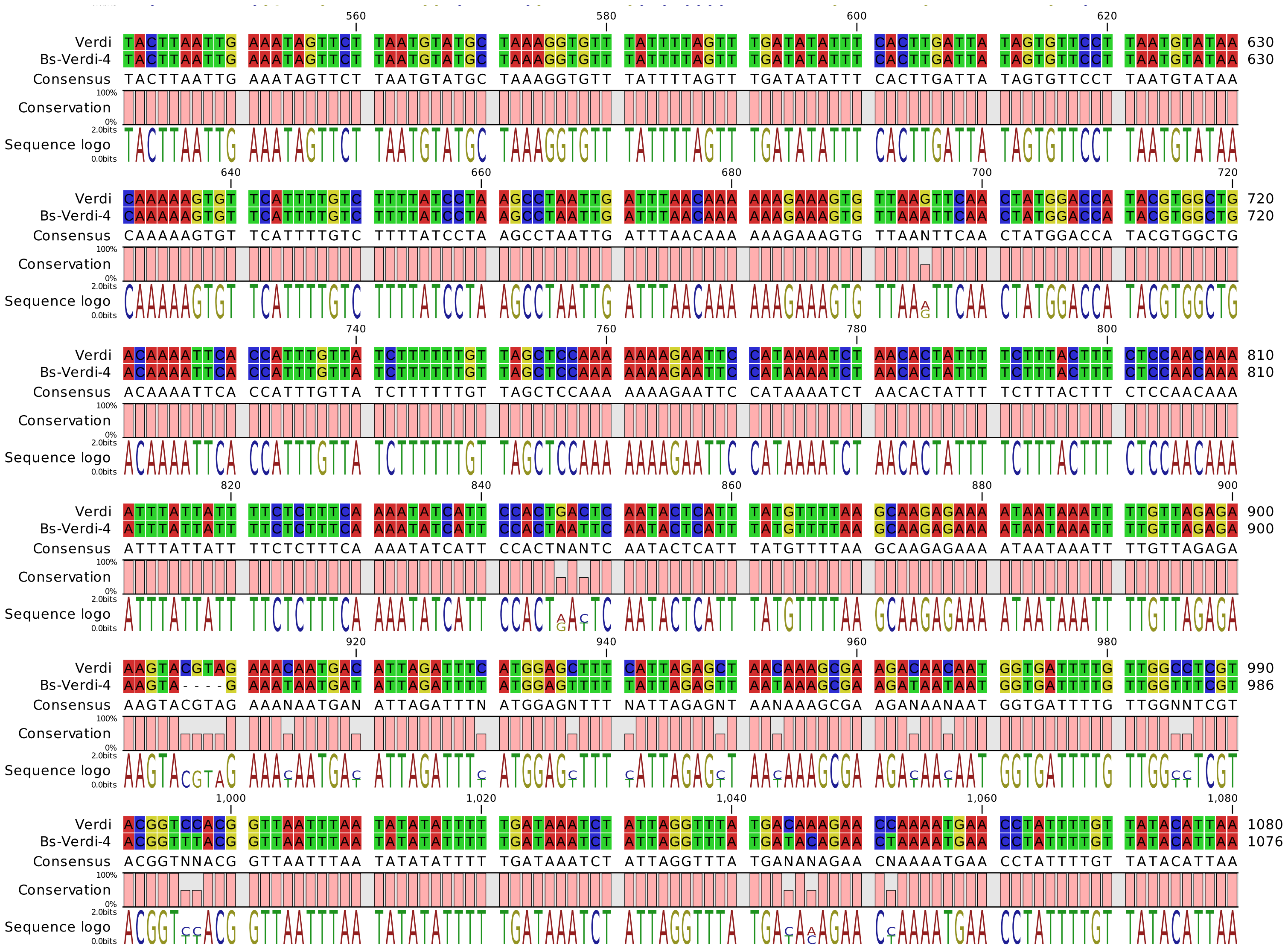
Verdi TACTTAATTG AAATAGTTCT TAATGTATGC TAAAGGTGTT TATTTTAGTT TGATATATTT CACTTGATTA TAGTGTTCCT TAATGTATAA 630
Bs-Verdi-4 TACTTAATTG AAATAGTTCT TAATGTATGC TAAAGGTGTT TATTTTAGTT TGATATATTT CACTTGATTA TAGTGTTCCT TAATGTATAA 630
Consensus TACTTAATTG AAATAGTTCT TAATGTATGC TAAAGGTGTT TATTTTAGTT TGATATATTT CACTTGATTA TAGTGTTCCT TAATGTATAA
Verdi CAAAAAGTGT TCATTTTGTC TTTTATCCTA AGCCTAATTG ATTTAACAAA AAAGAAAGTG TTAAGTTCAA CTATGGACCA TACGTGGCTG 720
Bs-Verdi-4 CAAAAAGTGT TCATTTTGTC TTTTATCCTA AGCCTAATTG ATTTAACAAA AAAGAAAGTG TTAAATTCAA CTATGGACCA TACGTGGCTG 720
Consensus CAAAAAGTGT TCATTTTGTC TTTTATCCTA AGCCTAATTG ATTTAACAAA AAAGAAAGTG TTAANTTCAA CTATGGACCA TACGTGGCTG
Verdi ACAAAATTCA CCATTTGTTA TCTTTTTTGT TAGCTCCAAA AAAAGAATTC CATAAAATCT AACACTATTT TCTTTACTTT CTCCAACAAA 810
Bs-Verdi-4 ACAAAATTCA CCATTTGTTA TCTTTTTTGT TAGCTCCAAA AAAAGAATTC CATAAAATCT AACACTATTT TCTTTACTTT CTCCAACAAA 810
Consensus ACAAAATTCA CCATTTGTTA TCTTTTTTGT TAGCTCCAAA AAAAGAATTC CATAAAATCT AACACTATTT TCTTTACTTT CTCCAACAAA
Verdi ATTTATTATT TTCTCTTTCA AAATATCATT CCACTGACTC AATACTCATT TATGTTTTAA GCAAGAGAAA ATAATAAATT TTGTTAGAGA 900
Bs-Verdi-4 ATTTATTATT TTCTCTTTCA AAATATCATT CCACTAATTC AATACTCATT TATGTTTTAA GCAAGAGAAA ATAATAAATT TTGTTAGAGA 900
Consensus ATTTATTATT TTCTCTTTCA AAATATCATT CCACTNANTC AATACTCATT TATGTTTTAA GCAAGAGAAA ATAATAAATT TTGTTAGAGA
Verdi AAGTACGTAG AAACAATGAC ATTAGATTTC ATGGAGCTTT CATTAGAGCT AACAAAGCGA AGACAACAAT GGTGATTTTG TTGGCCTCGT 990
Bs-Verdi-4 AAGTA----G AAATAATGAT ATTAGATTTT ATGGAGTTTT TATTAGAGTT AATAAAGCGA AGATAATAAT GGTGATTTTG TTGGTTTCGT 986
Consensus AAGTACGTAG AAANAATGAN ATTAGATTTN ATGGAGNTTT NATTAGAGNT AANAAAGCGA AGANAANAAT GGTGATTTTG TTGGNNTCGT
Verdi ACGGTCCACG GTTAATTTAA TATATATTTT TGATAAATCT ATTAGGTTTA TGACAAAGAA CCAAAATGAA CCTATTTTGT TATACATTAA 1080
Bs-Verdi-4 ACGGTTTACG GTTAATTTAA TATATATTTT TGATAAATCT ATTAGGTTTA TGATACAGAA CTAAAATGAA CCTATTTTGT TATACATTAA 1076
Consensus ACGGTNNACG GTTAATTTAA TATATATTTT TGATAAATCT ATTAGGTTTA TGANANAGAA CNAAAATGAA CCTATTTTGT TATACATTAA
Conservation
Sequence logo

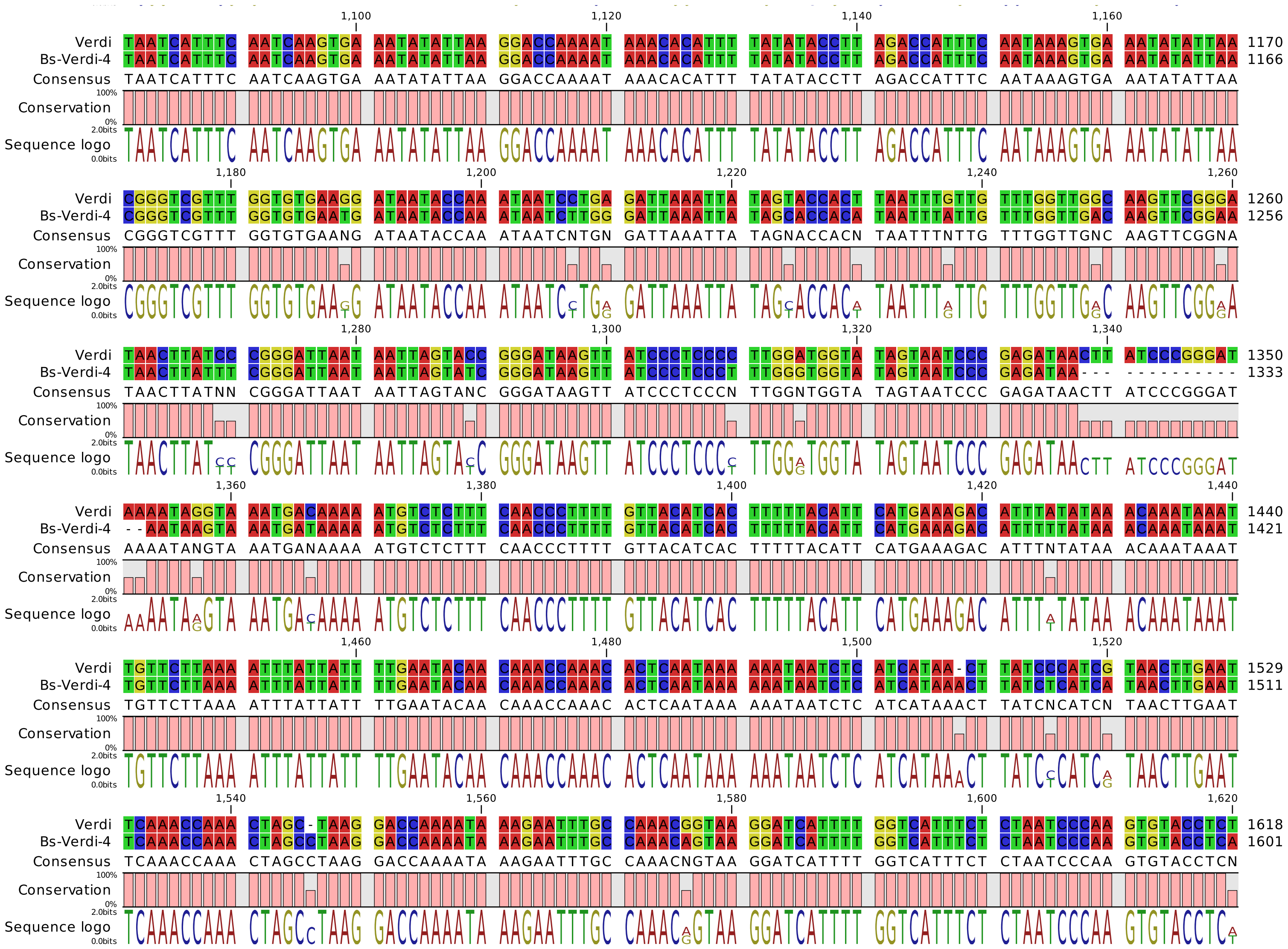
Verdi TAATCATTTC AATCAAGTGA AATATATTAA GGACCAAAAT AAACACATTT TATATACCTT AGACCATTTC AATAAAGTGA AATATATTAA 1170
Bs-Verdi-4 TAATCATTTC AATCAAGTGA AATATATTAA GGACCAAAAT AAACACATTT TATATACCTT AGACCATTTC AATAAAGTGA AATATATTAA 1166
Consensus TAATCATTTC AATCAAGTGA AATATATTAA GGACCAAAAT AAACACATTT TATATACCTT AGACCATTTC AATAAAGTGA AATATATTAA
Conservation
Sequence logo
Verdi CGGGTCGTTT GGTGTGAAGG ATAATACCAA ATAATCCTGA GATTAAATTA TAGTACCACT TAATTTGTTG TTTGGTTGGC AAGTTCGGGA 1260
Bs-Verdi-4 CGGGTCGTTT GGTGTGAATG ATAATACCAA ATAATCTTGG GATTAAATTA TAGCACCACA TAATTTATTG TTTGGTTGAC AAGTTCGGAA 1256
Consensus CGGGTCGTTT GGTGTGAANG ATAATACCAA ATAATCNTGN GATTAAATTA TAGNACCACN TAATTTNTTG TTTGGTTGNC AAGTTCGGNA
Conservation
Sequence logo
Verdi TAACTTATCC CGGGATTAAT AATTAGTACC GGGATAAGTT ATCCCTCCCC TTGGATGGTA TAGTAATCCC GAGATAACTT ATCCCGGGAT 1350
Bs-Verdi-4 TAACTTATTT CGGGATTAAT AATTAGTATC GGGATAAGTT ATCCCTCCCT TTGGGTGGTA TAGTAATCCC GAGATAA--- ---------- 1333
Consensus TAACTTATNN CGGGATTAAT AATTAGTANC GGGATAAGTT ATCCCTCCCN TTGGNTGGTA TAGTAATCCC GAGATAACTT ATCCCGGGAT
Conservation
Sequence logo
Verdi AAAATAGGTA AATGACAAAA ATGTCTCTTT CAACCCTTTT GTTACATCAC TTTTTACATT CATGAAAGAC ATTTATATAA ACAAATAAAT 1440
Bs-Verdi-4 --AATAAGTA AATGATAAAA ATGTCTCTTT CAACCCTTTT GTTACATCAC TTTTTACATT CATGAAAGAC ATTTTTATAA ACAAATAAAT 1421
Consensus AAAATANGTA AATGANAAAA ATGTCTCTTT CAACCCTTTT GTTACATCAC TTTTTACATT CATGAAAGAC ATTTNTATAA ACAAATAAAT
Conservation
Sequence logo
Verdi TGTTCTTAAA ATTTATTATT TTGAATACAA CAAACCAAAC ACTCAATAAA AAATAATCTC ATCATAA-CT TATCCCATCG TAACTTGAAT 1529
Bs-Verdi-4 TGTTCTTAAA ATTTATTATT TTGAATACAA CAAACCAAAC ACTCAATAAA AAATAATCTC ATCATAAACT TATCTCATCA TAACTTGAAT 1511
Consensus TGTTCTTAAA ATTTATTATT TTGAATACAA CAAACCAAAC ACTCAATAAA AAATAATCTC ATCATAAACT TATCNCATCN TAACTTGAAT
Conservation
Sequence logo
Verdi TCAAACCAAA CTAGC-TAAG GACCAAAATA AAGAATTTGC CAAACGGTAA GGATCATTTT GGTCATTTCT CTAATCCCAA GTGTACCTCT 1618
Bs-Verdi-4 TCAAACCAAA CTAGCCTAAG GACCAAAATA AAGAATTTGC CAAACAGTAA GGATCATTTT GGTCATTTCT CTAATCCCAA GTGTACCTCA 1601
Consensus TCAAACCAAA CTAGCCTAAG GACCAAAATA AAGAATTTGC CAAACNGTAA GGATCATTTT GGTCATTTCT CTAATCCCAA GTGTACCTCN
Conservation
Sequence logo

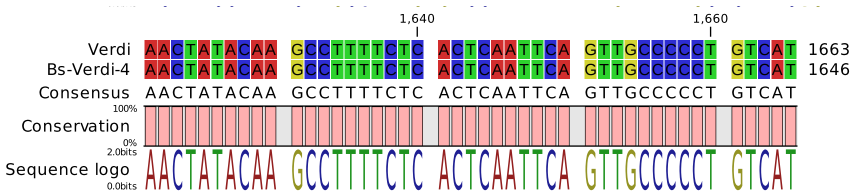
1,640
1,660
Verdi AACTATACAA GCCTTTTCTC ACTCAATTCA GTTGCCCCCT GTCAT 1663
Bs-Verdi-4 AACTATACAA GCCTTTTCTC ACTCAATTCA GTTGCCCCCT GTCAT 1646
Consensus AACTATACAA GCCTTTTCTC ACTCAATTCA GTTGCCCCCT GTCAT
100%
Conservation
0%
2.0bits
Sequence logo
0.0bits
